# *MECP2* duplication causes aberrant GABA pathways, circuits and behaviors in transgenic monkeys: neural mappings to patients with autism

**DOI:** 10.1101/728113

**Authors:** Dan-Chao Cai, Zhiwei Wang, Tingting Bo, Shengyao Yan, Yilin Liu, Zhaowen Liu, Kristina Zeljic, Xiaoyu Chen, Yafeng Zhan, Xiu Xu, Yasong Du, Yingwei Wang, Jing Cang, Guang-Zhong Wang, Jie Zhang, Qiang Sun, Zilong Qiu, Shengjin Ge, Zheng Ye, Zheng Wang

## Abstract

*MECP2 gain- and loss-of-function* in genetically-engineered monkeys demonstrably recapitulate typical phenotypes in patients, yet where *MECP2* mutation affects the monkey brain and whether/how it relates to autism pathology remains unknown. Using expression profiles of 13,888 genes in 182 macaque neocortical samples, we first show that *MECP2* coexpressed genes are enriched in GABA-related signaling pathways. We then perform analyses on multiple phenotypic levels including locomotive and cognitive behavior, resting-state electroencephalography and fMRI in *MECP2* overexpressed and wild-type macaque monkeys. Behaviorally, transgenic monkeys exhibit hyperactive and repetitive locomotion, greater separation anxiety response, and less flexibility in rule switching. Moreover, decreased neural synchronization at beta frequency (12-30 Hz) is associated with greater locomotion after peer separation. Further analysis of fMRI-derived connectomics reveals widespread hyper- and hypo-connectivity, where hyper-connectivity prominently involving prefrontal and cingulate networks accounts for deficits in cognitive flexibility. To map *MECP2*-related aberrant circuits of monkeys to the pathological circuits of autistic patients, individuals in a large public neuroimaging database of autism were clustered using community detection on functional connectivity patterns. In a stratified cohort of 49 autisms and 72 controls, the dysfunctional connectivity profile particularly in prefrontal and temporal networks is highly correlated with that of transgenic monkeys, as is further responsible for the severity of social communicative deficits in patients. Through establishing a circuit-based construct link between transgenic animal models and stratified clinical patients, the present findings with explicable biological causes are potentially amenable to translation for accurate diagnosis and evaluation of future treatments in autism-related disorders.

**One sentence summary:** We identify shared circuit-level abnormalities between *MECP2* transgenic monkeys and a stratified subgroup of human autism, and demonstrate the translational need of a multimodal approach to capture multifaceted effects triggered by a single genetic event in a genetically-engineered primate model.

## Introduction

Psychiatric disorders including autism spectrum disorder (ASD) are increasingly prevalent (*1*), and substantially heterogeneous in genetic bases and phenotypic architecture, consisting of subtypes with distinct biological mechanisms (*2, 3*). To achieve precise diagnosis and treatment, patient stratification based on biological measures has been encouraged in recent research initiatives (*3, 4*). Using non-invasive electrophysiological and neuroimaging measures, neural subtypes or biotypes of mental illnesses have been identified based on distinct patterns of brain network dysfunction, which are disease-relevant and predictive of treatment responsiveness (*5*). Recent efforts have also been made to bring genetic disorders with a high penetrance of ASD to the forefront of translational efforts to find treatments for subpopulations of mechanism-based classification of ASD (*6*). Although this strategy holds significant potential in filtering the complex etiology of autism, how specific genes with rare copy number variants contribute to functional alterations in brain circuitry, and what neural circuit basis may underlie particular pathological behaviors in ASD remains elusive (*6*).

An alternative route to dissect clinical heterogeneity and pursue the underlying neuropsychiatric mechanisms is genetic animal modeling, where certain causative genes in these biologically homogeneous samples are purposely manipulated to recapitulate some dimensions of core symptoms in psychiatric patients (*7*). One could expect that such animal models with a clear genetic basis would (at least partially) share circuit constructs with certain psychiatric neural subtypes, thereby providing valuable opportunities for probing the gene-circuit-behavior casual chain of events underlying complex brain disorders (*7, 8*). Here we focus on a transgenic macaque model of ASD with mutations in Methyl-CpG binding protein 2 (*MECP2*) (*9*). As extensively demonstrated in both humans (*10*) and rodent models (*11*), *MECP2* is one of few exceptional genes causing autistic features including intellectual disability, motor dysfunction, anxiety and social behavior deficits. In a previous study, we reported the successful application of lentiviral-mediated methods to produce genetically engineered macaque monkeys carrying extra copies of *MECP2* that manifested less active social contact and increased stereotypical behaviors (*9*). However, the relevant brain circuits that mediate the causal effects of atypical *MECP2* levels on phenotypic symptoms in transgenic monkeys remain essentially unexplored. In addition, it is unknown whether the neural circuit observed in the monkey model will map neatly onto homologs in specific neural subtypes of autism patients. To address these questions, we employ a combination of *MECP2*-coexpression network analysis, locomotive and cognitive behavioral tests (*12–14*), resting-state functional magnetic resonance imaging (rsfMRI) (*7, 15*) and electroencephalography (EEG) recordings (*16*), all of which are commonly administered in primate species and potentially amenable to cross-species translation into clinical diagnosis and future development of therapeutic interventions for autism-related disorders. We further conduct a cross-species circuit mapping of dysfunctional connectivity profiles between transgenic monkeys and subgroups of ASD patients who are stratified based on whole-brain resting-state connectivity patterns.

## Results

### Association of *MECP2* coexpression network with GABA function

To gain system-level insight into how *MECP2* mutations in transgenic (TG) monkeys affect the underlying biological process, we applied a weighted gene coexpression network analysis (WGCNA) to the gene expression data from the neocortex of adult macaque monkeys (*Macaca mulatta*). Expression profiles of 13,888 genes from 182 public samples were utilized to construct the gene coexpression network (see Supplementary Materials and Methods and Fig. 1A). We found that *MECP2* belonged to a module consisting of 159 coregulated genes and was positively correlated with the Module Eigengene (ME) (*r* = 0.66, *P* < 2.2×10^-16^, Fig. 1, B and C). Further functional enrichment analysis revealed that this module was highly associated (*P* < 0.01, FDR corrected) with four human diseases including unipolar depression (*P* = 0.002), four gene pathways including GABA (gamma-aminobutyric acid) function (*P* = 0.002), and three cellular components including clathrin-sculpted GABA vesicular transport (*P* = 0.002) (Fig. 1D). Next we explored the transcriptome of cortical samples from three deceased TG and four wild-type (WT) monkeys [PMID: 26808898] to further examine the role of this GABA-related module in *MECP2* transgenic monkeys. By comparing the expression abundance of *MECP2* top-linked genes between TG and WT monkeys, we found that most of the genes that exhibit the strongest association with *MECP2* expression were dramatically regulated in TG monkeys. Among those genes, 5 of them were significantly up-regulated (pink color in Fig. 1C) while 18 genes were significantly down-regulated (purple color in Fig. 1C). More details of these dysregulated genes are provided in fig. S1. Together the results in both TG and WT monkeys indicate a direct effect of *MECP2* alteration on the expression of those network adjacent genes.

**Fig. 1.**
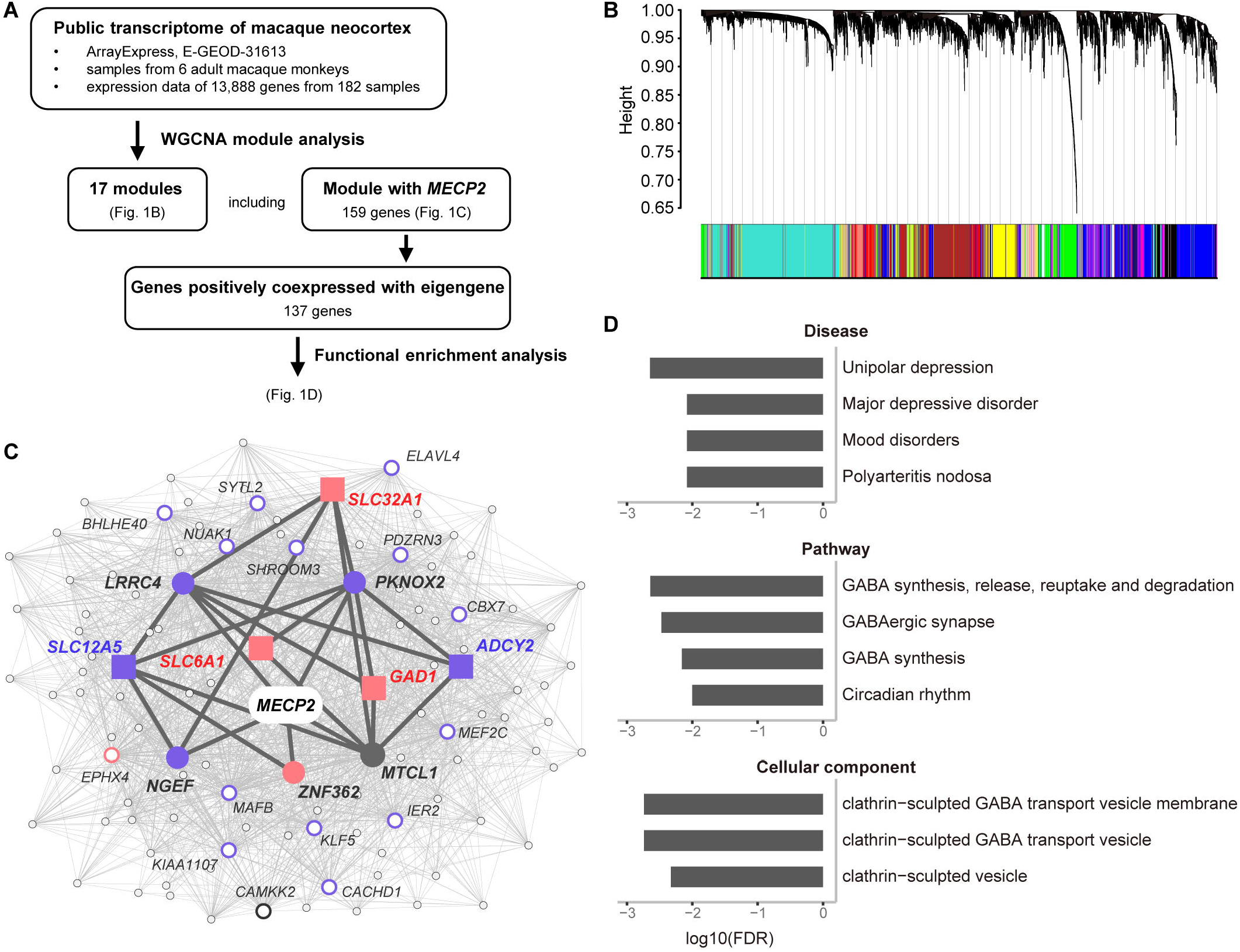
WGCNA and functional enrichment analysis. **(A)** Overview of weighted gene coexpression network analysis using public transcriptional data of 182 macaque cortical samples, and functional enrichment analysis on identified gene module coexpressed with *MECP2*. **(B)** Network analysis dendrograms representing assignment of 13,888 genes to 17 modules. **(C)** *MECP2* coexpression network. Directed neighbors of *MECP2* (*n* = 5) are represented with filled circles; genes enriched in GABA-related pathways (*n* = 5) are represented with filled squares; the top 20 genes with larger node degree are represented with empty circles; and the other second-order neighbors of *MECP2* are represented with smaller gray circles. Pink and purple colors indicate upregulation and downregulation in *MECP2* transgenic monkeys, respectively. Dark gray lines (edges) indicate the path from *MECP2* to GABA associated genes. **(D)** Four human diseases, four gene pathways, and three cellular components are significantly enriched in constrained genes (FDR-corrected *P* < 0.01). The enrichment *P*-values were calculated and corrected using ToppGene Suite.

### Abnormal locomotive behavior in transgenic monkeys

At 12∼18 months of age, these TG monkeys exhibited an increased frequency of repetitive circular locomotion (*9*). When they reached ∼55 months of age, we repeated the analysis on spontaneous locomotive behaviors in five TG and sixteen age-matched WT monkeys (*Macaca fascicularis*) (see Supplementary Materials and Methods). Repetitive locomotion including circular routing (movie S1), tumbling (movie S2), and cyclic routing in other more complex paths (movie S3) were observed. General locomotion time and repetitive index in home cage activity were analyzed by two independent observers to indicate the performance in locomotive and repetitive behavior, respectively. The inter-rater reliability of the two measures was 96.3% and 98.1%, respectively. The two measures were not correlated in either TG (*r* = −0.04, *P* = 0.952) or WT (*r* = −0.35, *P* = 0.182) monkeys. The total time spent in general locomotion for TG monkeys was significantly greater than for WT monkeys (*P* = 0.005, *Hedge’s g* = 1.55, two-sample Student’s *t*-test). The repetitive index of locomotion was also significantly higher in TG monkeys (*P* = 0.001, *Hedge’s g* = 1.94), as presented in Fig. 2B and table S1.

**Fig. 2.**
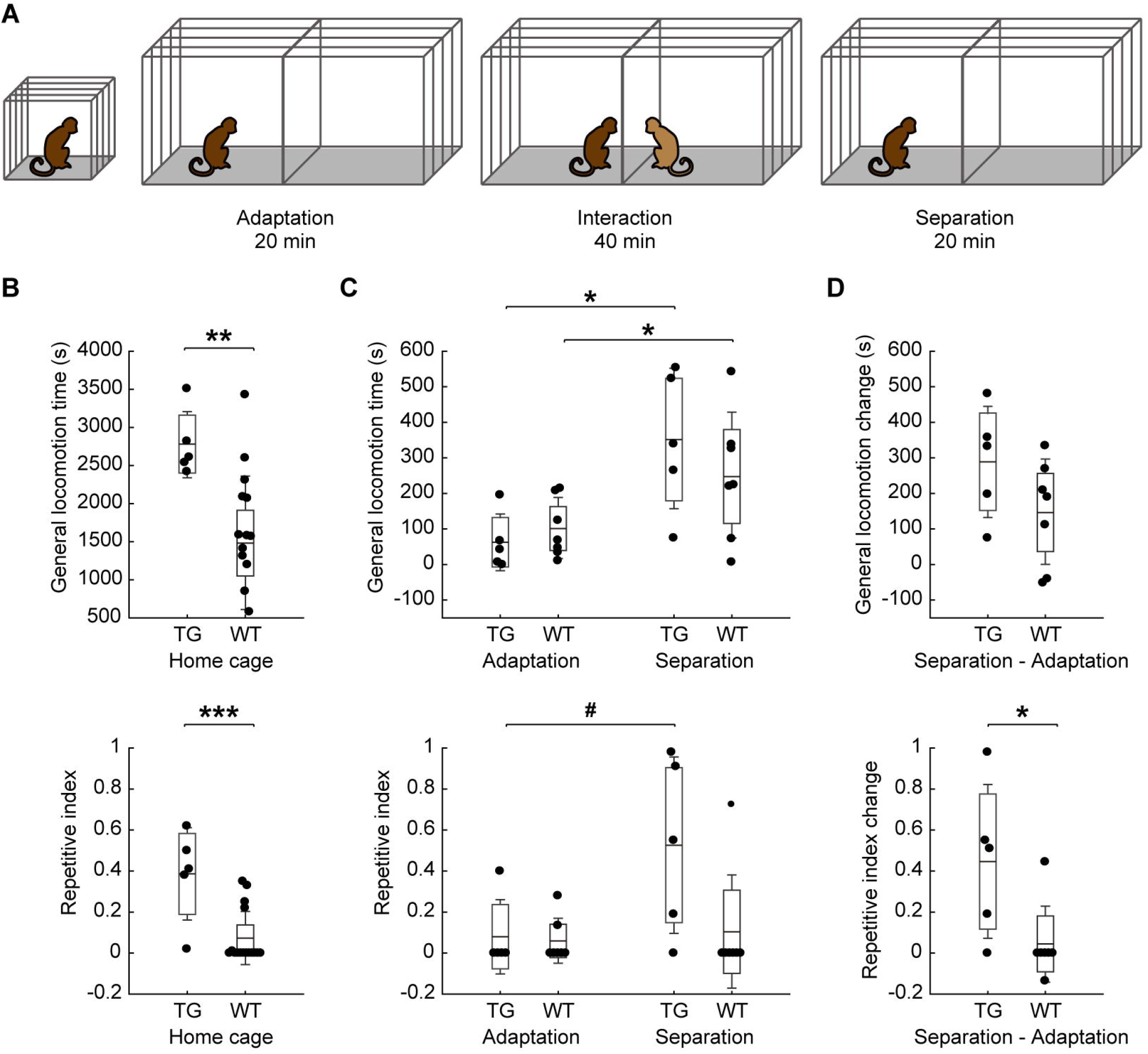
Excessive locomotive behaviors during home cage observation and peer separation test in TG monkeys. **(A)** Peer separation paradigm. The test monkey was first transferred from the home cage to a two-compartment test cage and stayed alone for 20 min. A peer monkey was then introduced into the other compartment for 40 min, during which the two monkeys were able to have visual, auditory and olfactory interactions. The test monkey stayed alone for another 20 min after the peer monkey was removed. Home cage observation and peer separation test were carried out on separate days. **(B)** Locomotion in home cage in terms of general locomotion time (upper panel) and repetitive index (lower panel). **(C)** Locomotion in the last 10 min of the adaptation and the first 10 min of the separation period in peer separation test. **(D)** Induced change in locomotive behaviors after peer separation. *** *P* < 0.0001, ** *P* < 0.001, * *P* < 0.05, # *P* < 0.1.

TG monkeys also exhibited increased anxiety at an early age in addition to excessive stereotyped locomotion (*9*). To further examine the relationship between abnormal locomotion and trait anxiety, we designed a novel test (Fig. 2A) based on a classical peer separation paradigm (*17*) (see Supplementary Materials and Methods). Absolute change of locomotive measures after separation with familiar peers was estimated to indicate the locomotive response to separation anxiety in five TG and seven WT monkeys. Both TG and WT monkeys showed increased locomotion time after peer separation compared to the adaptation period (*P* = 0.014; *P* = 0.040, respectively; Fig. 2C, upper panel). No group difference was found in the amount of change in locomotion time (*P* = 0.138, *Hedge’s g* = 0.87; Fig. 2D, upper panel). TG monkeys also showed a marginally significant increase in repetitive index (*P* = 0.056; Fig. 2C, lower panel) whereas WT monkeys showed no such change (*P* = 0.547). There was a significant group difference in the amount of change in repetitive index (*P* = 0.032, *Hedge’s g* = 1.34; Fig. 2D, lower panel). As a control study, the same test monkeys were further paired with unfamiliar peer monkeys to mediate the degree of separation anxiety. No group difference in induced behavioral changes was found in the case of unfamiliar peer separation in either general locomotion time (*P* = 0.643) or repetitive index (*P* = 0.953). More details of locomotion activity in the familiar and unfamiliar peer separation test are provided in table S2 and table S3, respectively.

### Increased regressive errors in reversal learning task

To probe potential cognitive inflexibility induced by *MECP2* dysfunction (*18*), we trained five TG and four WT monkeys to perform a discrimination and reversal learning task on a touch screen (*12–14*) (see Supplementary Materials and Methods). After the animals learned stimulus-directed touching (fig. S2A and movie S4), they were trained to discriminate between green and blue color, and to select the correct color for food reward when presented with two colored buttons (Fig. 3A and movie S5). The rewarded color was initially set to green, then reversed to blue after acquisition of the previous rule of color-reward association, and then randomly switched across sessions (Fig. 3, A and B). Animals had to discover the currently rewarded color through trial and error, and apply that rule in a session of 80 consecutive trials (Fig. 3B and movies S6-7).

**Fig. 3.**
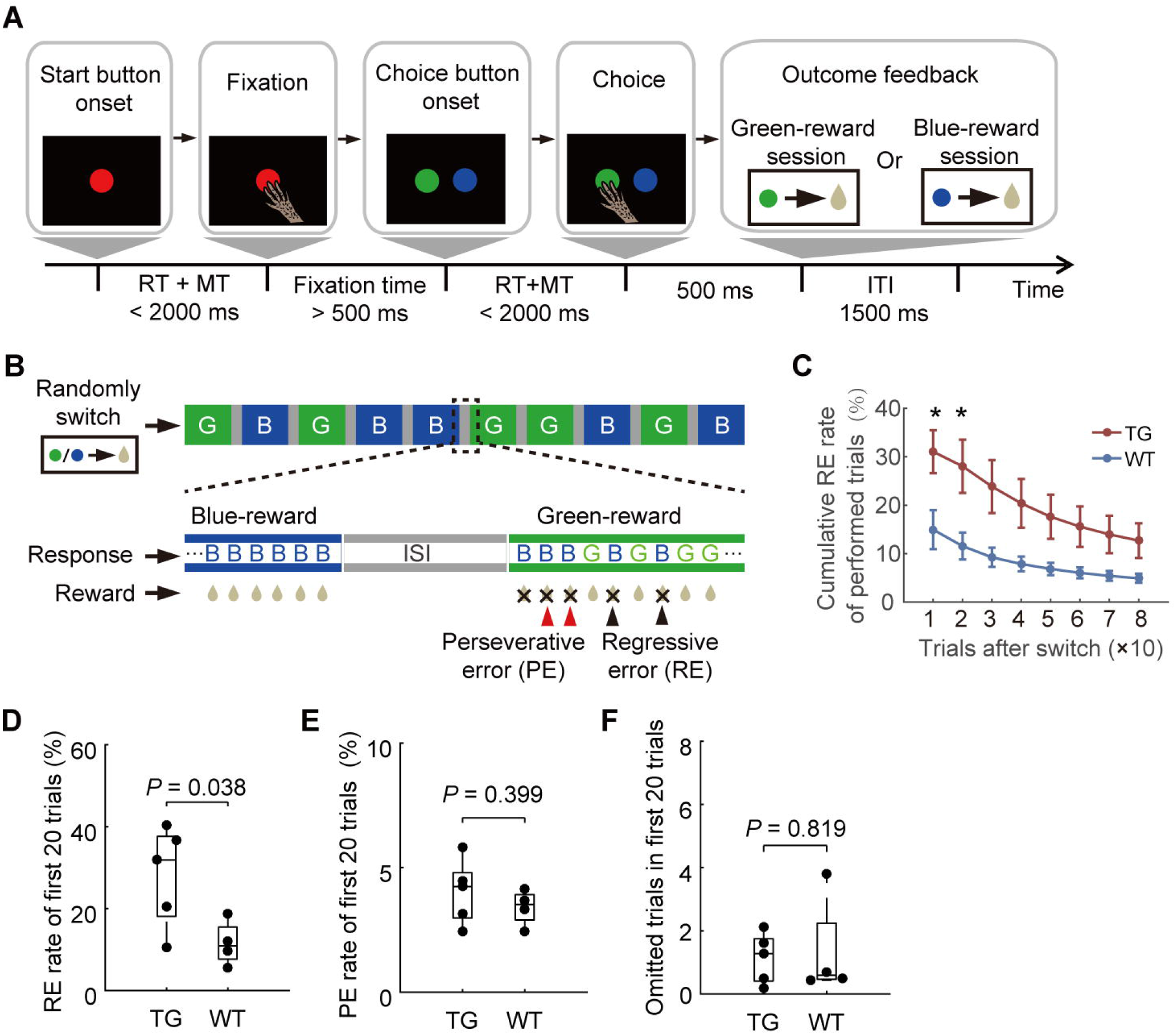
Cognitive flexibility tests in *MECP2* mutants and WT monkeys. **(A)** Temporal sequence of events in the color discrimination and reversal learning task. After subjects maintained a 500 ms hand fixation, two choice buttons (blue and green colors) were presented at randomized positions on the screen across trials. Subjects had to select the correct color within 2000 ms to get a 500 ms reward. RT, reaction time; MT, movement time; ITI, inter-trial interval. **(B)** An example of task sequence in the reversal learning test. The test included 10 sessions per day, 80 trials per session. The color assigned for reward was pseudo-randomly switched between sessions. The inter-session interval (ISI) was about 1 min. Red and black arrows indicate perseverative errors and regressive errors, respectively. G, green-reward session; B, blue-reward session. **(C)** The cumulative regressive error rate of performed trials after rule switch was analyzed (*n* = 5, TG; *n* = 4, WT; \**P* < 0.05, two-sample Student’s *t*-test). Error bars denote standard error of mean. **(D-F)** The percentage of regressive errors (D), perseverative errors (E) and number of omitted trials (F) in TG and WT groups in first 20 performed trials after rule switch (Student’s *t*-test).

Two types of error after rule switches were assessed. Perseverative errors were trials in which monkeys continued to select the previously rewarded color following negative feedback but prior to the first correct response, suggesting a failure to switch to a new rule. By contrast, regressive errors were trials in which monkeys continued to select the previously rewarded color after the first correct response, thus representing a preference for the old rule or an inability to maintain a new rule (Fig. 3B). Notably, TG monkeys exhibited a significantly greater regressive error rate for the first twenty performed trials than WT monkeys (Fig. 3C and D, *P* = 0.038, *Hedge’s g* = 1.47, two-sample Student’s *t*-test), while they did not demonstrate an increased rate of perseverative errors (*P* = 0.443, *Hedges’ g* = 0.46, Fig. 3E). Meanwhile, TG and WT monkeys omitted a similar number of trials (*P* = 0.710, *Hedges’ g* = 0.26, Fig. 3F), and performed equivalently on acquiring both stimulus-directed touching (fig. S2B) and stimulus-reward association rules (fig. S2, E to G). Individual performance at all training stages is presented in fig. S2 (B to D, H to L).

### Decreased beta synchronization in transgenic monkeys

We conducted multi-channel scalp EEG recordings in five TG and sixteen WT monkeys (see table S4 for more details). Light anesthesia state was maintained during data collection by experienced anesthesiologists. Details on EEG data acquisition and analysis, and anesthesia maintenance are provided in Supplementary Materials and Methods. Fig. 4A and Fig. 4B show the spatial location of electrodes (see table S5 for full description) in a three-dimensional reconstruction of individual anatomical MRI scan and a standard macaque brain template (*19*), respectively. Spectral power at each recording site, as well as phase synchronization strength between each pair of recording sites were estimated and compared across two groups in six canonical frequency bands (*20*), namely delta (1–4 Hz), theta (4–8 Hz), alpha (8–12 Hz), beta (12–30 Hz), low gamma (30–60 Hz), and high gamma (60–100 Hz). No significant group difference between TG and WT monkeys was found in the relative power of either frequency band after correction for multiple comparisons (fig. S3).

**Fig. 4.**
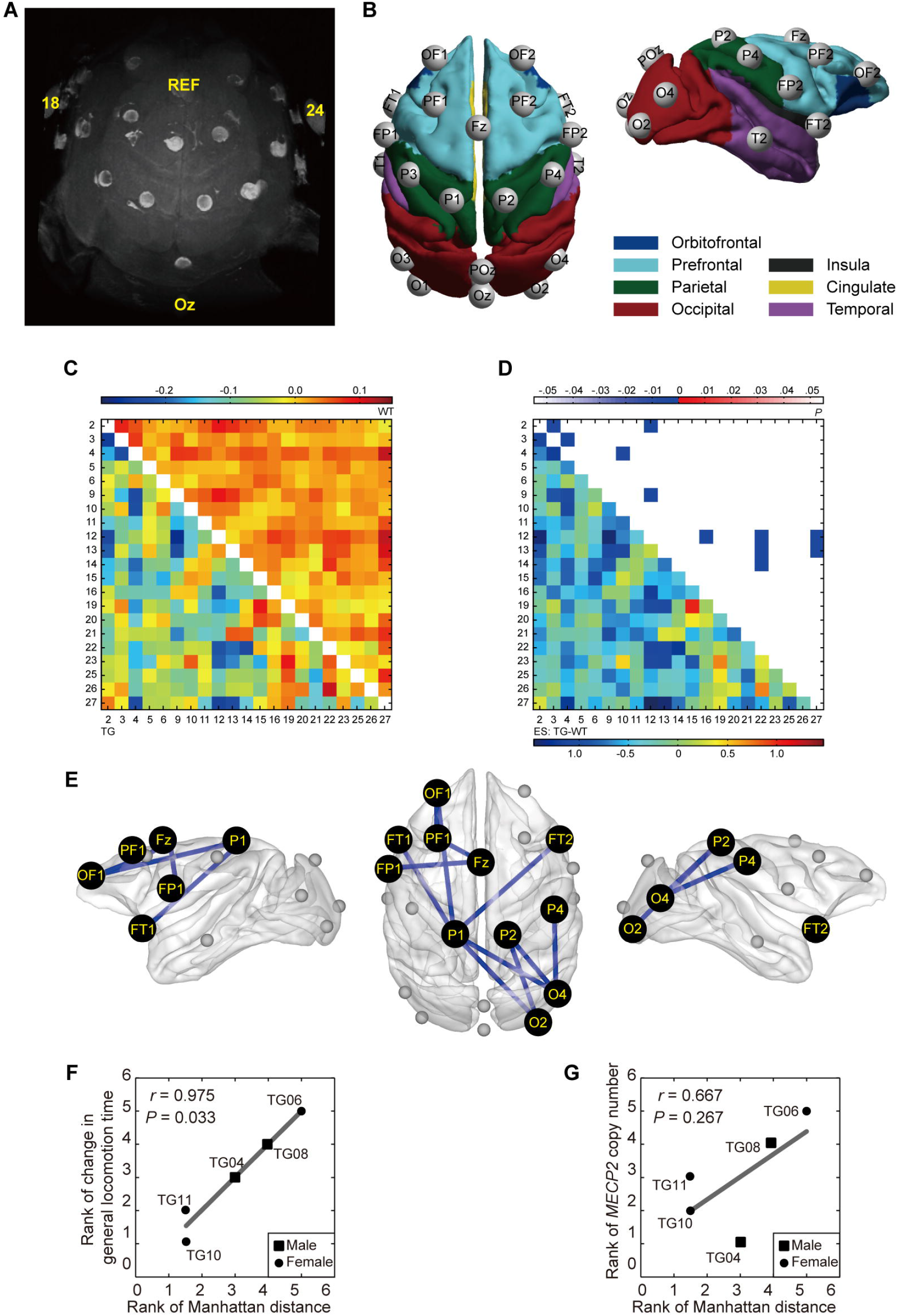
Decreased beta (12-30 Hz) synchronization in fronto-parieto-occipital networks in TG monkeys. **(A)** Three-dimensional reconstruction of monkey TG04 wearing the EEG cap based on T2-weighted MRI images. Bright circles indicate electrode rings. Bright ellipses above circles indicate marked electrodes, including the reference (REF), most posterior (Oz) and most lateral (18/24) electrodes. The ground electrode (not shown here) was located far from the cortex beneath the left eye. Six recording channels far from the cortex (including 18/24) were disabled for subsequent recording. **(B)** Spatial organization of 21 active electrodes after mapping to the cortical surfaces of the F99 template brain. Brain lobes were displayed in different colors. The electrodes were named after the spatial location. See table S5 for more details. **(C)** Averaged covariates-free neural synchronization networks of TG (bottom-left) and WT (top-right) monkeys. The numbers indicate the electrode ID as in table S5. The color bar denotes the logarithm of dwPLI. **(D)** Effect sizes (ESs, *Hedge’s g* value, bottom-left) and statistical *P* values (top-right) of statistical comparison between TG and WT groups. Eleven connections widely distributed in fronto-parieto-occipital networks were identified significantly decreased in TG monkeys (edge-wise *P* < 0.005, cluster-level corrected *P* = 0.010, more details in fig. S3 and table S6). **(E)** Topographic distribution of abnormal connections was illustrated in the F99 template brain. **(F-G)** Association of the overall abnormality in beta synchronization (Manhattan distance) with general locomotion time change after peer separation (F) and *MECP2* copy number (G). Associations were evaluated via Spearman’s rank correlation for TG monkeys only.

Inter-site phase synchronization was estimated via de-biased weighted phase-lag index (dwPLI) (*21*). Significant group differences were found specifically in the beta band at the network level. Group-averaged beta synchronization networks of TG and WT monkeys with age (linear and quadratic) and gender effect regressed out are presented in Fig. 4C, together with the corresponding effect size (*Hedges’ g* value) of individual neural synchronization network connections (Fig. 4D). We identified a fronto-parieto-occipital network that showed a statistically significant decrease in beta synchronization in TG monkeys (Fig. 4E, edge-wise *P* < 0.005, cluster-level corrected *P* = 0.010). This network consisted of two connections within the frontal lobe (OF1-PF1, PF1-Fz) and four fronto-parietal connections (Fz-FP1, OF1-P1, FT1-P1, FT2-P1) in the left hemisphere, and five parieto-occipital connections (P1-O2, P1-O4, P2-O2, P2-O4, P4-O4) in the right hemisphere (more details are specified in fig. S4 and table S6). The averaged neural synchronization matrices of all frequency bands are shown in fig. S5. In contrast, no significant group difference in neural synchronization of other frequency bands was observed.

Given the strong evidence of copy-number-dependent symptom severity observed in both animal models and patients (*10, 11*), we attempted to explore potential associations among genetic variability, circuit abnormality and behavioral phenotypes in the current experimental setting. We adopted Manhattan distance to quantitatively describe the overall abnormal extent of the connectivity fingerprint of individual TG relative to WT controls (*15, 22*). Such a ‘distance measure’ can determine whether different fingerprints are ‘close’ or ‘far’ from each other. Manhattan distances based on connections with abnormal beta synchronization were calculated for each TG monkey. Spearman’s correlation analysis was performed in a pairwise manner to probe associations among gene (*MECP2* copy number), circuit (Manhattan distance based on abnormal beta synchronization), and behavior (general locomotion time and repetitive index in home cage, locomotion change induced by familiar peer separation, and error rate in reversal learning task) in five TG monkeys. The overall abnormality in beta synchronization was significantly correlated with locomotion time change (*r* = 0.975, *P* = 0.033, Fig. 4F) after familiar peer separation, but not with the *MECP2* copy number (*r* = 0.667, *P* = 0.267, Fig. 4G).

### Aberrant functional connectivity in transgenic monkeys

We applied a between-group comparison of functional connectome derived from intrinsic resting-state fMRI signals, which are highly sensitive and stable, and correlate well with brain gene expression (*23, 24*). Details on MRI data acquisition and analysis are provided in Supplementary Materials and Methods. We acquired whole-brain fMRI data and constructed connectivity networks with 94 parcellated brain regions (also termed nodes, see table S7 for a complete list of anatomical labels) in sixteen anesthetized macaques including five TG and eleven WT monkeys (table S8). Group-averaged connectivity networks of TG and WT monkeys with age and gender as covariates are subjected to two-sample Student’s *t*-tests, as shown in Fig. 5A. We identified a total of 50 functional connections that showed significant differences between TG and WT groups (edge-wise *P* < 0.001, cluster-level corrected *P* = 0.019, top-right triangle in Fig. 5B), and then quantified the extent of these group effects by using corresponding effect size (ES) (bottom-left triangle in Fig. 5B). Connectivity strength of these connections in all monkeys is presented in fig. S6. Out of 94 nodes, 40 widespread throughout the entire brain were found containing abnormal connections with remarkably large effect sizes (table S9). The degree of abnormality of each node, i.e., the sum of effect sizes of all abnormal edges connecting to this node (*ESn*), is presented in the interlayer of Fig. 5C. The right primary somatosensory cortex (S1) showed decreased connectivity with large *ESn* (top 25%, blue bars in Fig. 5C). In contrast, increased connectivity regions with the top 25% largest *ESn* (red bars) covered a wide range of frontal cortex including left polar and right centrolateral prefrontal cortex (PFCpol, PFCcl), right lateral orbitofrontal cortex (PFCol) and frontal eye field (FEF), cingulate areas including bilateral retrosplenial and left anterior cingulate cortices (CCr and CCa), and other areas including right medial parietal (PCm) and left parahippocampus (PHC). The distribution of altered functional connections in brain lobes are summarized in Fig. 5D (*Z* scores in bottom-left triangle, see Supplementary Materials and Methods), after the non-uniform parcellation of the brain template was accounted for in a standardized residual analysis (*25*). The distribution was preferentially biased to prefrontal and cingulate cortices (*P* < 0.05, Bonferroni correction, top-right triangle in Fig. 5D).

**Fig. 5.**
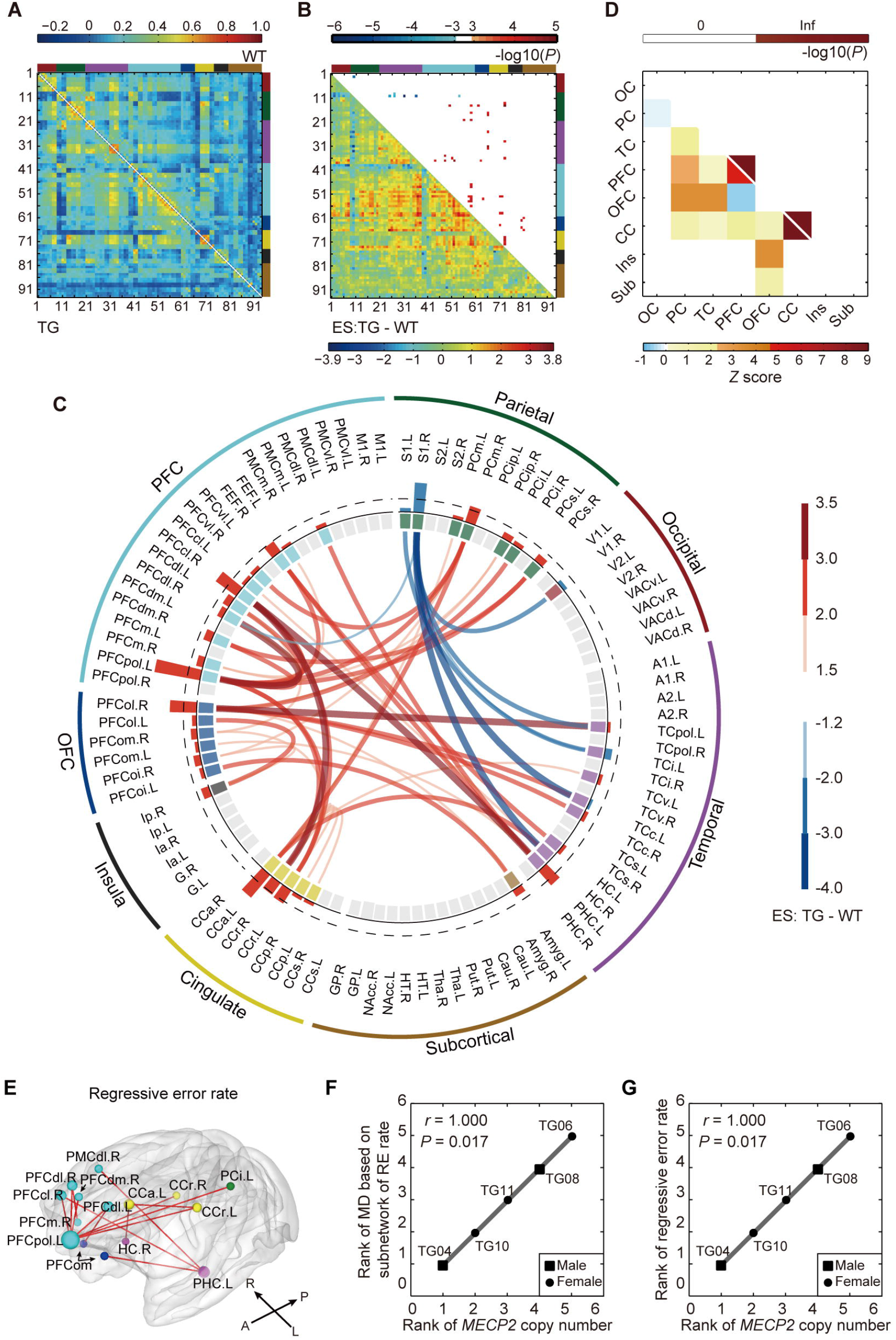
Disrupted neural circuits in transgenic monkeys. **(A)** Covariate-free connectivity network matrices for TG (bottom-left) and WT (top-right) monkeys. **(B)** Effect sizes (ESs) of TG versus WT (bottom-left) are shown with corresponding *P* values (top-right, cluster-level corrected *P* < 0.05, and edge-wise *P* < 0.001). Brain nodes are sorted and organized according to the regions/lobes as listed in table S7. **(C)** Disrupted functional connections and corresponding brain nodes in TG monkeys compared with WT. The red and blue bars in the interlayer indicate positive and negative ES of each brain node (*ESn*), respectively, which was defined by the sum of the ESs of all abnormal connections to this node. The dashed line labels the top 25% of absolute *ESn*. The abbreviations and parcellation of brain nodes are listed in table S7. **(D)** Spatial distribution of disrupted connections across the brain is plotted (bottom-left) and the regions with statistical significance are highlighted (top-right, *P* < 0.05, Bonferroni correction). Inf, infinite; OC, occipital cortex; PC, parietal cortex; TC, temporal cortex; PFC, prefrontal cortex; OFC, orbitofrontal cortex; CC, cingulate cortex; Ins, insula; Sub, subcortical areas. **(E)** Subnetwork associated with regressive error rate in reversal learning task at the significance level of *P* < 0.05 after correction for multiple comparisons. **(F)** Gene-circuit association between *MECP2* copy number and circuit abnormality underlying regressive error rate. MD, Manhattan distance; RE, regressive error. **(G)** Gene-behavior association between *MECP2* copy number and regressive error rate. Associations were evaluated via Spearman’s rank correlation for TG monkeys only.

The unique functional connectivity profile of individual TG monkeys was calculated in a similar way to their neural connectivity profiles. We plotted the connectivity fingerprint of each TG, in which topological alterations in individual nodes are clearly illustrated (fig. S7). We found that Manhattan distances based on abnormal functional connections of monkeys TG04, TG06, TG08, TG10 and TG11 are 8.06, 9.64, 9.17, 6.98 and 9.17, respectively, all of which except TG10 were statistically different from WT monkeys (*P* = 0.025, 0.000, 0.000, 0.155 and 0.000, permutation test, fig. S7). The Manhattan distance was marginally significantly associated with regressive error rate in cognitive flexibility test (*r* = 0.872, *P* = 0.067). To further dissect the neural circuit that may underlie a dimensional phenotype in the identified network abnormalities, we quantitatively evaluated the relation between each abnormal edge and behavioral measures, and then adjusted the statistical significance at the cluster-level (see Supplementary Materials and Methods). Fourteen functional connections that mostly stemmed from left PFCpol and left PHC were correlated with the regressive error rate in the reversal learning task (edge-wise *P* < 0.05, cluster-level corrected *P* = 0.006, Fig. 5E). Strikingly, *MECP2* copy number in TG monkeys was significantly associated with the Manhattan distance of these dysfunctional connections (*r* = 1.000, *P* = 0.017, Fig. 5F) and with the regressive error rate (*r* = 1.000, *P* = 0.017, Fig.5G). Seven other functional connections that mostly emanated from right S1 were correlated with general locomotion time in home cage (edge-wise *P* < 0.01, cluster-level corrected *P* = 0.005 and *P* = 0.037, respectively). However, the overall abnormality in this subnetwork was not associated with *MECP2* copy number (*r* = 0.500, *P* = 0.450). More details on behavior-related edges are provided in table S10.

### Pathological circuits and symptoms in patients with autism

We proceeded to analyze the functional connectivity networks of human subjects using the same strategy as in the monkey experiments (Fig. 6, A and B). Given the substantial heterogeneity and large spectrum of clinical symptoms among ASD patients (*26*), individuals were first clustered into subgroups with relatively homogenous brain network organization (Fig. 6C). Pathological circuits regulating behavioral phenotypes in different autism subtypes were then evaluated via case-control comparison (Fig. 6D) and symptom correlation analysis (Fig. 6E).

**Fig. 6.**
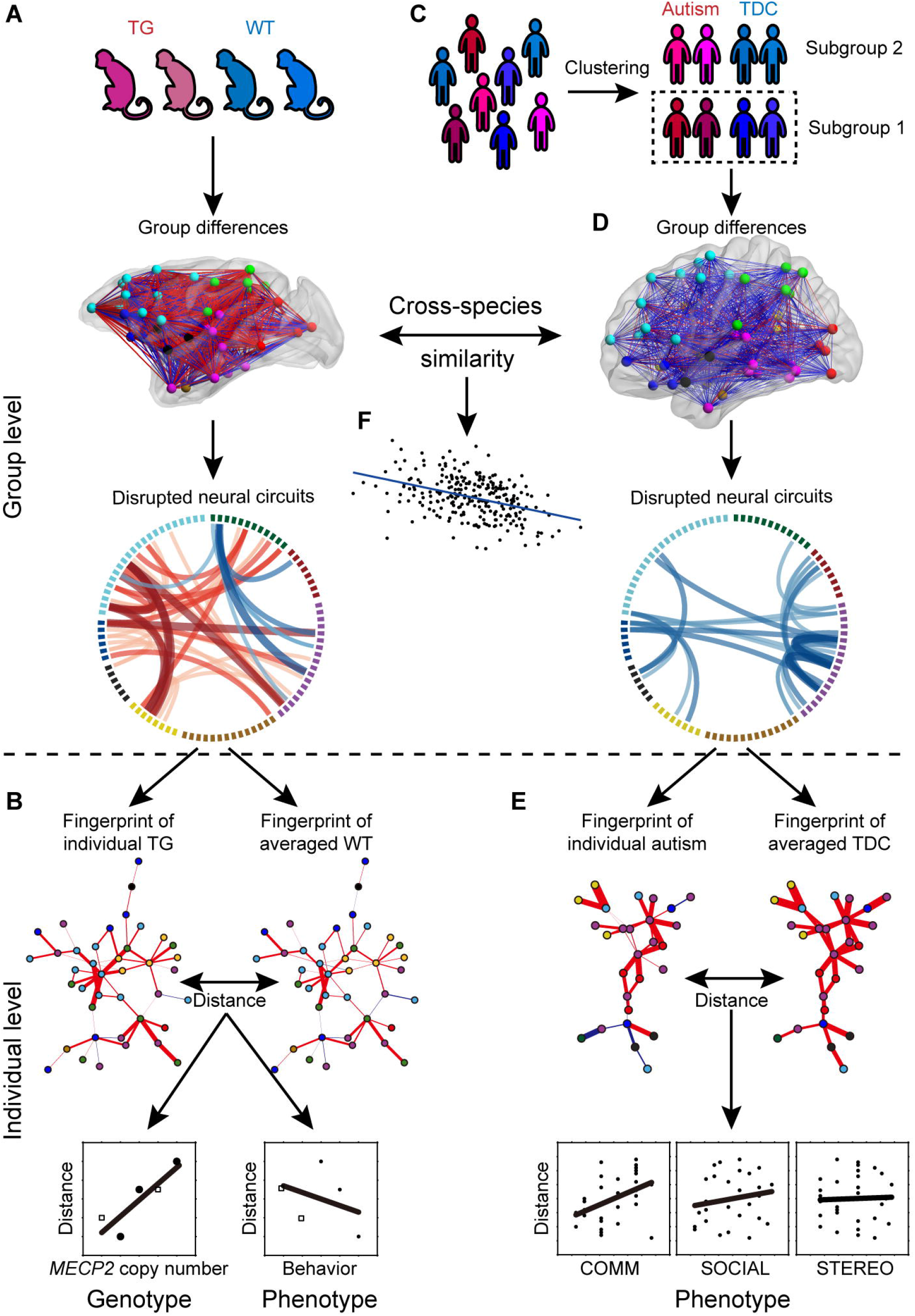
Schematic of comparative analysis strategy between monkeys and humans. **(A)** Group-level differences of functional connectivity networks between transgenic (TG) and wild-type (WT) monkeys were statistically compared and their effect sizes were evaluated. **(B)** The connectivity fingerprint of each TG monkey was identified and evaluated by Manhattan distance (MD) relative to an averaged WT monkey. The relationships between genetic, circuitry and behavioral aberrations were explored. **(C)** Stratification of human participants was done through data-driven clustering based on the resting-state fMRI connectivity network. **(D)** Group-level differences of connectivity networks between autism and typically developing controls (TDC) were analyzed for each subgroup using the same strategy as in the monkey experiments. **(E)** Connectivity fingerprint of individual autism was identified and evaluated by MD. Their relations to the severity of dimensional symptoms assessed by clinical scores were explored. **(F)** Cross-species similarity of altered functional homologs in TG monkeys and autistic patients was estimated based on (A) and (D).

We used public data from the Autism Brain Imaging Data Exchange (ABIDE), which shares MRI scans and phenotypic information from 539 patients with ASD and 573 typically developing controls (TDCs) (*27*). After basic demographic and diagnostic screening, 90 adolescents with clinical diagnoses of autism and 140 matched TDCs were enrolled (table S11). Covariate-free (including age, gender, full-scale IQ, and scanning sites) connectivity networks were then reconstructed for these subjects based on the same brain parcellation scheme used for monkeys (table S7). The enrolled subjects were then clustered into subgroups based on their rsfMRI connectivity network using community detection on the inter-participant spatial correlation matrix (see Supplementary Materials and Methods). Two clusters were derived: subgroup 1 consisted of 49 autism and 72 TDCs, and subgroup 2 consisted of 41 autism and 68 TDCs (fig. S8A, tables S12 and S13). Brain network homogeneity within each subgroup was significantly improved for both autism and TDCs (*P* < 0.05, fig. S8B). No discernable differences in phenotypic and demographic characteristic were found between these two subgroups of autism (fig. S8C).

As in the monkey dataset, we repeated the same statistical analysis for both subgroups in parallel (Fig. 7 and fig. S9). In subgroup 1, 32 significant hypo-connections were observed in 27 brain nodes (edge-wise *P* < 0.001, cluster-level corrected *P* = 0.026, Fig. 7, B and C, and table S14), most of which were located between temporal cortex (TC) and PFC, between TC and orbitofrontal cortex (OFC), between TC and occipital cortex, and within TC (*P* < 0.05, Bonferroni correction, Fig. 7D). Regions with the top 25% largest *ESn* were right PFCpol, right PFCol, bilateral ventral TC (TCv), right TC polar (TCpol), right superior TC (TCs) and right primary auditory cortex (A1) (blue bars of interlayer in Fig. 7C). In contrast, no abnormal connections in subgroup 2 withstood correction for multiple comparisons (fig. S9).

**Fig. 7.**
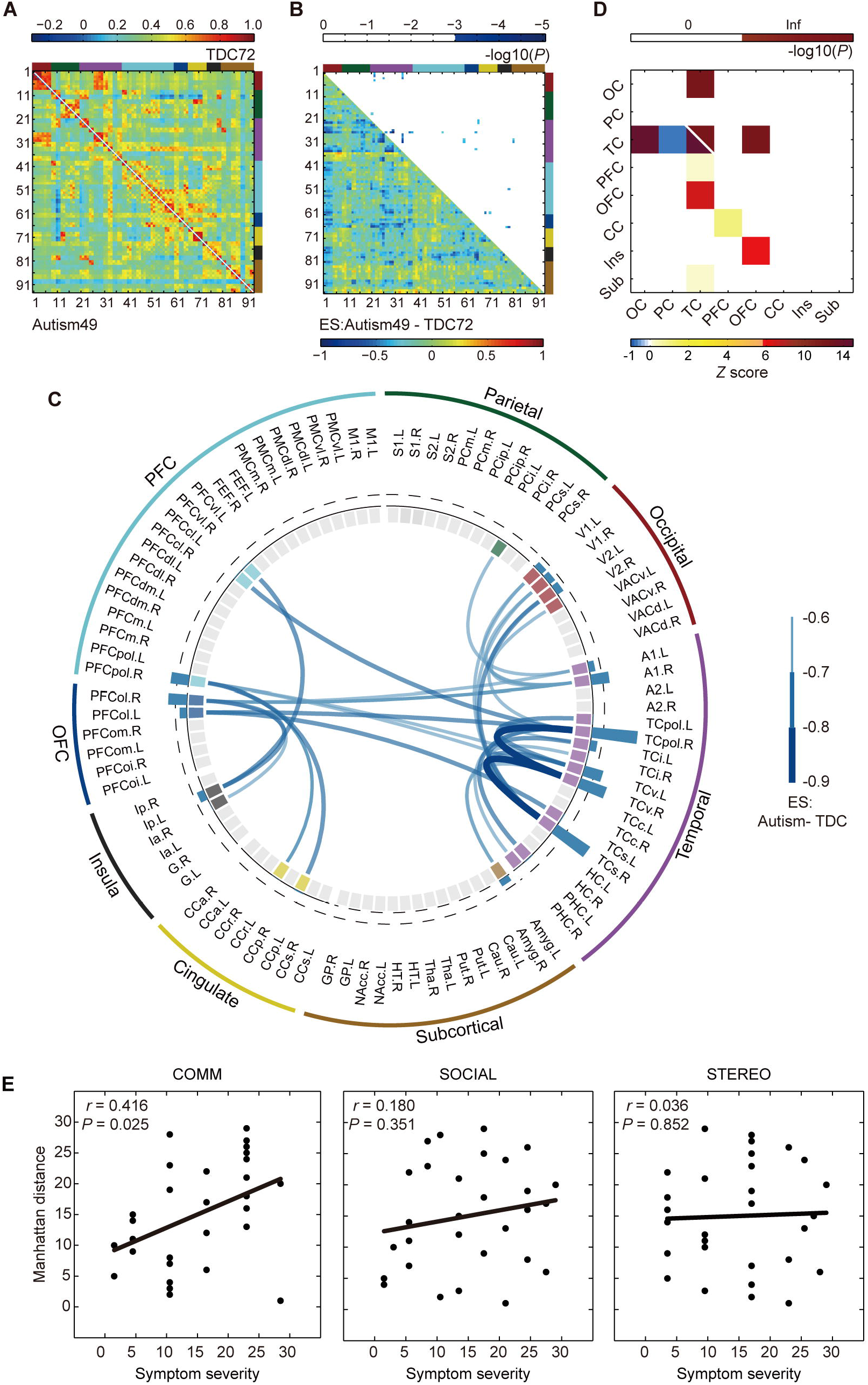
Disrupted neural circuits in a subgroup of patients with autism. **(A)** Covariate-free connectivity network matrices for autism (bottom-left) and TDC (top-right) in subgroup 1. **(B)** Effect sizes (ESs) of autism versus TDC (bottom-left) are shown with corresponding *P* values (top-right, cluster-level *P* < 0.05, edge-wise *P* < 0.001). The brain nodes are sorted and organized according to the regions/lobes as listed in table S7. **(C)** Disrupted functional connections and corresponding brain nodes in patients with autism compared with TDC. The blue bar in the interlayer indicates negative *ESn*, which was defined as the sum of the ESs of all abnormal connections to this node. The dashed line labels the top 25% of absolute *ESn*. The abbreviations and parcellation of brain nodes are listed in table S7. **(D)** Spatial distribution of disrupted connections across the brain is plotted (bottom-left) and highlighted with statistical significance (top-right, *P* < 0.05, Bonferroni correction). Inf, infinite; OC, occipital cortex; PC, parietal cortex; TC, temporal cortex; PFC, prefrontal cortex; OFC, orbitofrontal cortex; CC, cingulate cortex; Ins, insula; Sub, subcortical areas. **(E)** Associations between Manhattan distances of each autism patient and the severity of dimensional symptoms in communication (left), social interaction (middle), and stereotyped behaviors (right), respectively.

To map different dimensions/domains of autistic symptoms (as assessed by the Autism Diagnostic Observation Schedule, ADOS) onto the connectivity fingerprint in subgroup 1, we applied Manhattan distance to quantitatively describe the overall extent of circuit abnormality in individual autism relative to an averaged TDC (*22, 26*) (see Supplementary Materials and Methods). In 29 of 49 participants with research-reliable ADOS scores, we observed a significant correlation between communication scores and their Manhattan distances (Spearman’s *r* = 0.416, *P* = 0.025), but not with social interaction or restricted, repetitive behavior (Fig. 7E and table S15). No subnetworks underlying specific symptoms were further derived from the disrupted network.

### Cross-species comparison of connectivity fingerprints

After independently evaluating functional abnormalities in monkey and human networks, we next asked whether any relationship existed between the two (Fig. 6F). Although no abnormal connections were shared between transgenic monkeys and subgroup 1 of patients with autism (Figs. 5C and 7C), there were overlaps in abnormal brain nodes (Fig. 8A), which include lateral prefrontal areas (left PFCcl, right PFCvl), lateral orbitofrontal areas (bilateral PFCol), left temporal areas (TCi, TCv and TCpol), left cingulate (CCp and CCr), left PHC, left amygdala, right inferior parietal area, and right primary visual area (V1). Comparing to this subset of patients, monkeys exhibited further abnormality in other subdivisions of PFC (bilateral PFCdl, left PFCpol, and right PFCcl, PFCdm and PFCm) and OFC (bilateral PFCoi and PFCom), FEF, central TC (TCc), PCm, S1, and hippocampus. Abnormal brain areas specific to the patient subgroup include bilateral superior and right medial TC (TCs and TCm), bilateral insula, bilateral A1, and left V1 and bilateral secondary visual areas (V2).

**Fig. 8.**
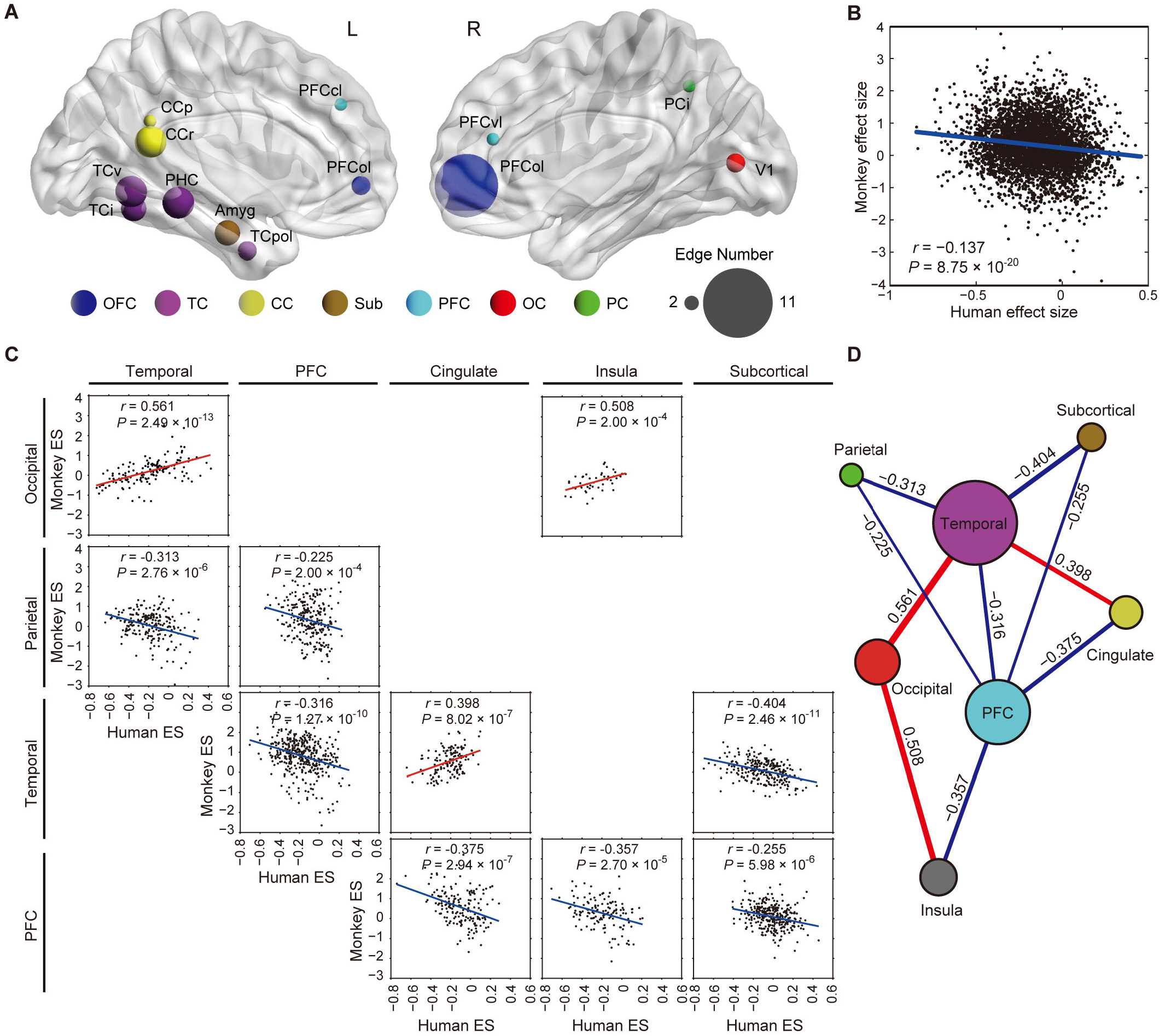
Cross-species comparison of connectivity fingerprints in monkeys and humans. **(A)** Abnormal brain regions shared between transgenic monkeys and human patients. The size of nodes indicates the total number of abnormal edges connected to the node in both species. OFC, orbitofrontal cortex; TC, temporal cortex; CC, cingulate cortex; Sub, subcortical areas; PFC, prefrontal cortex; OC, occipital cortex; PC, parietal cortex. **(B)** Spatial correlation between the entire effect size (ES) matrices of TG versus WT monkeys (shown in Fig. 5B) and patients with autism versus TDC in subgroup 1 (shown in Fig. 7B). (**C)** Spatial correlation of ES matrices in different brain lobes between monkeys and humans. Each scatter plot displays statistically significant correlations in different lobes between the two primates (*P* < 0.05, Bonferroni correction), as summarized in **(D)**. The size of brain lobes in the spider plot indicates the sum of all significant correlation coefficients of ESs between the lobe and all other lobes of the two primates.

We further conducted cross-species comparison at network level by calculating pairwise spatial correlations between group-different monkey networks and group-different human networks (lower triangle of effect size matrices in Figs. 5B and 7B, see Supplementary Materials and Methods). The connectivity fingerprint of monkeys substantially correlated only with human subgroup 1 (*r* = −0.137, *P* = 8.75×10^-20^, Fig. 8B), but not with subgroup 2 (*r* = 0.013, *P* = 0.408, fig. S10). To further test whether such cross-species similarity was uniform across different parts of the brain, we divided the whole brain circuitry into eight lobes/regions and computed their corresponding spatial correlations. As plotted in Fig. 8C, correspondence of connectivity fingerprints between PFC and temporal, parietal, cingulate, insula and subcortical areas, and between temporal and parietal, cingulate, occipital and subcortical areas were remarkably high between the two primates (*P* < 0.05, Bonferroni correction, table S16). Taking all between-lobe/regions spatial correlations into account, we found that homologous networks between TG monkeys and autistic patients largely center on PFC and temporal lobes, both of which show strong cross-species correlation with five other lobes/regions (Fig. 8D).

## Discussion

### *MECP2*-related GABA dysfunction and beta desynchronization

The frequency-dependent finding in electrophysiological activity likely reflects *MECP2*-induced dysfunction in specific molecular pathways. Evidence from transgenic rodents has demonstrated that *MECP2* dysfunction alters synchrony and the overall excitation/inhibition balance in brain circuits (*28*), with greater influence on GABAergic neurons (*29*). The present transcriptome and functional enrichment analysis of macaque neocortex reveals a strong association between *MECP2* coexpressed genes and GABA function, implying altered GABAergic action in the mutant monkeys. Moreover, results from both intracellular recording and genetic association studies suggest that generation of cortical beta oscillation depends on GABAergic neurotransmission (*30, 31*). As such, we propose that the disrupted beta synchronization observed in transgenic monkeys is very likely a consequence of the *MECP2*-induced malfunction of GABAergic neurons. Similar dysfunction of GABAergic signaling (*32*) and reduction in beta synchronization during resting state (*33*) have been reported in ASD patients. Although no associations between spontaneous beta coherence and behavioral symptoms have been reported in human patients thus far, this may be a potential diagnostic biomarker for autism which merits future validation.

### Circuit and behavior disruptions in transgenic monkeys and autistic humans

The prevailing hypothesis of disrupted cortical connectivity, in which deficiencies in the way the brain coordinates and synchronizes activity among different regions account for clinical symptoms of ASD, is central to the etiology and accurate diagnosis of ASD (*26, 34–36*). However, human heterogeneity in terms of complex genetic backgrounds, diverse clinical comorbidities, different genders, various developmental trajectories and medication status is a general issue and formidable challenge, causing diverse and contradictory EEG (*35, 37, 38*) and imaging findings of ASD (*26, 27, 34*). Therefore, in this study, we first stratified all enrolled subjects through a data-driven approach to improve the inter-participant homogeneity in whole-brain connectivity patterns. Disease-related disruptions in brain networks were then assessed via case-control comparison in subgroups (*39*). As a result, a hypo-connected network mainly located in prefrontal and temporal areas was identified in one subgroup. Several brain areas underlying language processing and verbal communication were found disrupted in this autism subgroup, including primary auditory and visual areas, and superior and medial temporal areas (*40*). The biological significance of the identified circuit was supported by the association between its overall disruption and communication deficits in the autism sample.

Cognitive flexibility deficits are proposed as potential psychological constructs underpinning major autistic domains, including restricted and repetitive behavior, atypical social interaction and abnormal communication (*18, 41*). In remarkable consistency with our observation in monkeys, ASD patients are impaired at maintaining newly-rewarded responses and inclined to revert back to previously-reinforced choices, which partially explains the increased severity of restricted and repetitive behaviors (*42*). Interestingly, *MECP2* copy number positively correlates with the increased rate of regressive errors, suggesting a gene dosage-dependent severity of the phenotype in these monkeys similar to that observed in human autism (*10*). Our circuit-behavior analysis further pins down a pertinent distributed network involving medial, orbital and lateral PFC regions, premotor, anterior and rostral cingulate, inferior parietal, hippocampal and parahippocampal areas, responsible for their performance in the reversal task. This finding is compatible with recent fMRI results in ASD patients performing a modified reversal learning task, where reduced activation was found in frontal and parietal networks that support flexible choices (*43*). Future studies are needed to investigate the role of cognitive flexibility in autistic etiology, especially its link to social communication and repetitive behavior.

### Cross-species neural mappings

We proposed a new cross-species comparison strategy (as illustrated in Fig. 6) involving human stratification and neural mappings from monkeys to human subgroups. Patient stratification before cross-species mapping is an important prerequisite because within the umbrella term “ASD” there exist very different subtypes of patients in this ABIDE large repository, meanwhile the ‘animal model’ is not expected to resemble the human disorder in every respect (*7*). As described above, data-driven stratification whereby multiple subgroups are biologically defined, should reveal multiple networks in parallel that may underlie different symptomatic domains in autism (*39*). Fortunately, we found that one of two derived subgroups demonstrated substantial overlapping abnormal brain regions with the monkey model (Fig. 8A), among which the lateral orbitofrontal cortex plays a vital role in disrupted circuits in both species (Fig. 5C and Fig. 7C).

It is somewhat surprising to observe a lack of shared connections between transgenic monkeys and autistic patients, even though the dysfunctional connectivity profiles of the two species were significantly correlated. As a matter of fact, it remains essentially unknown how exactly the brain circuits of primate species correspond to each other, due to evolutionary effects (*7, 22*). Given the substantial variations in functional connections subserving species-specific behavioral and cognitive adaptations (*7, 44*), topological characteristics of brain regions are more likely to be evolutionarily conserved. This assumption is supported as cross-species similarity in dysconnectivity profiles was more prominent in prefrontal and temporal cortices, thus implying etiological convergence between the monkey model and autism patients. Nevertheless, it is rather intriguing that shared abnormalities in disconnected circuits are tightly associated with cognitive flexibility deficits in monkeys whereas communication deficits in humans, respectively. This “bent” circuit-behavioral mapping at two ends may indicate a novel mechanism underlying social communication deficits in autism, where a specific neurocognitive defect – preference for old cognitive rules or failure to maintain new cognitive rules– disturbs executive function (*41*).

## Conclusions

We present the first neurophysiological and neuroimaging evidence in genetically-engineered macaque monkeys that genetic variants in *MECP2* may predispose individuals to abnormal locomotive and cognitive behaviors through dysregulation of neural and functional connectivity with prefrontal circuits. *MECP2* duplication-induced effect on neural connectivity of the primate model is both temporally (beta frequency range, 12-30 Hz) and spatially (fronto-parieto-occipital network, prefrontal and cingulate network) dependent. More importantly, dysfunctional connectivity profiles induced by *MECP2* duplication are mapped onto a subgroup of autistic individuals sharing similar profiles of brain circuits. Taken together, all present examinations in nonhuman primates that are conveniently adapted to human subjects not only hold crucial implications for accurate diagnosis of autism-related disorders, but also offer new insights into the development of targeted behavioral interventions that can improve atypical locomotion and social communication in autism-related disorders.

## Supplementary Materials

Materials and Methods

Supplementary References

Fig. S1. Heatmap for the expression of *MECP2* top correlated genes in the cortical regions of *MECP2* transgenic monkeys.

Fig. S2. Behavior results of color discrimination and reversal learning task.

Fig. S3. Topography of relative power differences between TG and WT monkeys for six frequency bands.

Fig. S4. Strength of connections with significantly decreased beta synchronization in TG monkeys.

Fig. S5. Averaged neural synchronization matrices for both TG and WT monkeys for six frequency bands.

Fig. S6. Scatter plot of abnormal connections in transgenic monkeys.

Fig. S7. Connectivity fingerprint of individual transgenic monkeys.

Fig. S8. Parsing heterogeneity in clinical cohorts.

Fig. S9. Abnormal functional connections of human autism in subgroup 2.

Fig. S10. Spatial correlation between entire effect size (ES) matrices of TG versus WT monkeys and human autism versus TDC in subgroup 2.

Table S1. *MECP2* copy number and locomotion of 5 TG and 16 WT monkeys in home cage.

Table S2. Locomotion of 5 TG and 7 WT monkeys in peer separation test with familiar peers.

Table S3. Locomotion of 5 TG and 7 WT monkeys in peer separation test with unfamiliar peers.

Table S4. EEG data collection from 5 TG and 16 WT monkeys.

Table S5. Spatial location and anatomical label of all recording electrodes.

Table S6. Connections with decreased beta synchronization in TG monkeys.

Table S7. Cortical and subcortical parcellation and abbreviations.

Table S8. MRI data collection from 5 TG and 11 WT monkeys.

Table S9. Abnormal functional connections in transgenic monkeys.

Table S10. Group different edges correlating with behavioral measures.

Table S11. Characteristics of all human participants.

Table S12. Characteristics of human participants in subgroup 1.

Table S13. Characteristics of human participants in subgroup 2.

Table S14. Disrupted functional connections in human autism of subgroup 1.

Table S15. Correlations between connectivity fingerprint of human autism in subgroup 1 and symptom severity.

Table S16. Correlations of group differences in connectivity networks between monkeys and humans (subgroup 1).

Movie S1. Example of repetitive circular routing behaviors.

Movie S2. Example of repetitive tumbling behaviors.

Movie S3. Example of repetitive cyclic routing in the path of a figure eight.

Movie S4. An example of monkey (TG04) performing stimulus-directed touching task.

Movie S5. An example of monkey (TG04) performing discrimination task.

Movie S6. An example of monkey (TG08) performing reversal learning task.

Movie S7. An example of monkey (WT032) performing reversal learning task.

## Acknowledgments

We would like to thank Hu Zhang, Ganxian Wang, Qinying Jiang and Wenwen Yu for their assistance to monkey training and data acquisition, and also thank Drs. Trevor Robbins, Freund Tamas, Charles Schroeder, Anna Roe, Ravi Menon, Stefan Everling and Muming Poo for their stimulating discussions and suggestions during the preparation of this study.

## Author contribution

D.-C.C., Z.-W.W., S.-Y.Y., and X.C., with the help of Y.W., J.C. and S.G., conducted the EEG experiments. D.-C.C., with the help of Z.W., analyzed the EEG data. Z.-W.W., T.-T.B., D.-C.C., and S.-Y.Y., with the help of J.C. and Z.W., conducted animal MRI experiments. Z.-W.W., with the help of Y.-F.Z. and Z.W., analyzed animal MRI data. T.-T.B., Y.-L.L. and S.-Y.Y., with the help of Z.Q., K.Z., Z.Y. and Z.W., designed animal behavioral experiments. T.-T.B. and Y.-L.L. conducted the reversal learning task and analyzed the behavioral data. S.-Y.Y. conducted the home cage observation and peer separation test. S.-Y.Y. and D.-C.C. analyzed the locomotive behaviors. Z.-W.W., with the help of D.-C.C., Y.-F.Z., X.X., G.-Z.W., Z.Y., Y.D. and Z.W., analyzed the human data and cross-species comparison. Z.L., with the help of J.Z., conducted the gene coexpression and enrichment analysis. Z.Q. and Q.S. generated the transgenic monkeys. X.X., Y.D., and G.-Z.W. contributed to data interpretation. Z.W., Z.Y., and S.G. conceived and supervised the project. Z.W. wrote the manuscript with the help of Z.Y., S.G., D.-C.C., Z.-W.W., and K.Z. All authors read and approved the manuscript.

