## Supplementary Materials for "*MECP2* duplication causes aberrant GABA pathways, circuits and behaviors in transgenic monkeys: neural mappings to patients with autism"

**This PDF file includes:**

Materials and Methods

Supplementary References

Fig. S1 to S10

Table S1 to S16

### Materials and Methods

#### Participants

All experimental procedures for nonhuman primate research in this study were approved by the Institutional Animal Care and Use Committee at the Institute of Neuroscience and the Biomedical Research Ethics Committee, Shanghai Institutes for Biological Sciences, Chinese Academy of Sciences, and conformed to National Institutes of Health guidelines for the humane care and use of laboratory animals.

Five *MECP2*-overexpressed transgenic (TG) monkeys (*Macaca fascicularis;* 3 female) and twenty wild-type (WT) monkeys (*Macaca fascicularis;* 11 female) participated in the present study. All TG monkeys participated in all three behavioral tasks. Subgroups of sixteen, seven and four WT monkeys participated in home cage observation, peer separation test and reversal learning task, respectively. All TG (age 5.60 ± 0.08 years, weight 3.90 ± 1.47 kg) and sixteen WT (age 5.66 ± 0.60 years, weight 5.51 ± 2.30 kg; 7 female) monkeys participated in the EEG experiment. All TG (age 4.40 ± 0.29 years, weight 3.26 ± 0.75 kg) and eleven WT (age 4.68 ± 0.46 years, weight 3.97 ± 1.36 kg; 7 female) monkeys participated in the MRI experiment.

#### Weighted gene coexpression network analysis (WGCNA)

We obtained gene expression data for the whole neocortex of six adult rhesus macaques from ArrayExpress (E-GEOD-31613) (*1*) and used the WGCNA package in R to construct the coexpression network of *MECP2* in the macaque brain. The experimental samples for mRNA profiling include two male and two female profiles for each of 10 cortical areas. We performed quality control processing for the raw expression data as described in previous work (*2*), including exclusion of outlier samples with inter-array correlations (IAC) < 2 standard deviations from the mean IAC, and correction for the cross-batch with the ComBat package.

Furthermore, we obtained the expression value of genes from their corresponding probes and selected the probe with the highest variation across samples to represent this gene. Finally, we included expression profiles of 13,888 genes in 182 samples in the following analysis.

The WGCNA algorithm employing dynamic cut-tree method was used to produce gene coexpression clusters (*3*). Seventeen clusters of genes, or modules, were identified in the dataset. We found that *MECP2* belonged to a module consisting of 159 genes and was positively correlated with the Module Eigengene (ME), defined as the first principle component of the module ($r = 0.64$, $P < 2.2\times10^{-16}$). Furthermore, using the 137 genes in the module that were also positively correlated with its ME, we performed functional enrichment analysis (using the Toppgene suit: https://toppgene.cchmc.org/) to infer the biological process to which the genes coexpressed with *MECP2* may be related.

**Monkey behavioral tasks**

**Locomotive behavior in home cage**

Spontaneous locomotive behaviors of five TG and sixteen WT monkeys in home cage were videotaped and analyzed. Monkeys were individually caged (length 0.9 m, width 0.8 m, height 0.9 m) in a room with ten to twelve roommates and monitored continuously. For each monkey, uninterrupted recordings were filmed for 20 min daily between 14:40 and 15:00 on four separate days. Behaviors of repetitive circular and general locomotion were analyzed by two independent trained observers, who were blinded to the genotypes of the monkeys. Inter-observer reliability was evaluated. Repetitive circular locomotion was defined as circular routing (movie S1), tumbling (movie S2), and cyclic routing in other more complex paths, e.g., the path of a figure eight (movie S3), for more than three cycles. General locomotion was defined as any form of locomotion occurring during non-resting states, with resting state referring to staying still for more than 3 s without obvious body movement (e.g., standing, sitting, and grooming). We further defined a repetitive index by dividing repetitive locomotion time by general locomotion time and used it as an indicator of repetitive behavior. The association between repetitive index and general locomotion time was evaluated via Pearson correlation in each group, respectively. Group differences in general locomotion time and repetitive index were statistically tested using independent two-sample *t*-tests.

**Locomotive behavior in peer separation test**

Pairs consisting of a test monkey and a peer monkey were used in the paradigm (Fig. 2A). Five TG and seven WT monkeys participated as the test monkey. Each of them was paired with two familiar and two unfamiliar WT monkeys on four separate sessions. Familiar peer monkeys had been housed in the same room as the test monkey for more than one year at the time of the experiment, while unfamiliar peer monkeys had never been housed in the same room as the test monkey. Eight WT monkeys participated as the familiar peer monkey, including seven WT monkeys who also participated as the test monkey. Four additional WT monkeys participated as the unfamiliar peer monkey.

The test apparatus was a two-compartment cage (length 3.4 m, width 1.9 m, height 1.7 m) made of steel rods. The two compartments were separated by electrochromic switchable glass, which changes from opaque to transparent when a voltage is applied. The entire cage was continuously monitored. Testing was carried out midafternoon from 15:00 to 17:30, and consisted of three periods, namely, adaption, interaction and separation. In the adaption period, the test monkey was transferred from home cage to the testing compartment, and stayed alone for 20 min. During the interaction period, the peer monkey was transferred to the other compartment. Behaviors of both monkeys were recorded for 40 min after the arrival of the peer monkey. The two monkeys were able to have visual, auditory and olfactory contact during this period. In the separation period, the peer monkey was removed from the cage before the test monkey was observed for another 20 min. The glass between the two compartments was transparent throughout the whole procedure except when the peer monkey was removed. The glass was switched to opaque temporally so that the test monkey would not be distracted by the food lure given to the peer monkey. Each test monkey was paired with at least two peer monkeys and tested on separate days.

Behavioral change induced by peer separation was estimated as the absolute change upon peer separation as compared to the baseline, the latter part of the adaptation period. Recordings from the last 10 min of the adaptation period and the first 10 min of the separation period were analyzed for locomotive behaviors. The same procedure was applied as to the home cage activity. For each test monkey, locomotion measures (general locomotion time and repetitive index) from different paired sessions were averaged. Thus, the data consisted of behavioral changes in two measures from all monkeys. The significance of behavioral changes across two periods was estimated via a paired two-sample *t*-test for each group. Group differences in induced locomotion changes were statistically tested via independent two sample *t*-test.

**Color discrimination and reversal learning task**

We employed a color discrimination and reversal learning paradigm to assess the TG monkeys' cognitive flexibility deficits. Cognitive flexibility entails that discrimination learning builds on stimulus-reward associations that are flexible in response to changing situational demands, which has been widely proposed to be pathognomonic of ASD. Five TG monkeys and four age-matched WT monkeys participated in this touchscreen-based psychophysical experiment. All monkeys were experimentally naïve and provided with a standard primate diet except on behavioral test days, on which 60-70 g of food pellets was delivered during morning and afternoon feeding, respectively, and 200 g fresh fruits at noon. The subjects were tested relatively unrestrained in behavioral primate chairs without head fixation. Before the experiment, animals were initially trained to habituate to pole and collar handling and placement in chairs. During experiments, each monkey was seated comfortably in a primate chair at a certain distance from the center of a frontally-positioned 19 inch touch screen (Elo 1991L open-frame LCD touch screen), which was determined by the length of each monkey's arm. The monkey’s right arm could freely move to touch the screen. The experimental task was programmed using the MonkeyLogic toolbox (*4*). The reward during the experiment for each monkey was a mixed liquid of 150 g fresh apple, 50 g banana, 60 g puree, 200 mL purified water and 45 g daily fodder, which was accurately supplied by a peristaltic pump (Longer Pump, BT100-1F) depending on the performance of each trial. The whole period of behavioral tests was videotaped.

The behavioral tests were conducted on weekdays. All monkeys were fasted 24 hours before the test day. To fully assess cognitive flexibility in these cynomolgus monkeys, we set up a reversal learning task with three different stages. Thus, we could maximize the number of subjects by utilizing a fairly simple learning task that many animals might be able to perform. For the first two stages and the beginning phase of stage 3, animals had to complete 4 sessions per day, each session containing 200 trials, and their performance was evaluated at each stage.

Stage 1: Stimulus-directed touching. Each trial began with presentation of a red fixation circle in the center of the screen with a field angle of 2 degrees. The animal had to learn to touch the screen within 2 s to receive a reward. Animals were first taught to touch the screen by manually being held by the hand to receive the reward. Once the animal could take the initiative to put the hand on the screen and sustain fixation on the screen, they were left alone to finish the task (movie S4). The longer the animals touched for, the greater the reward they would receive. The maximum reward time in each trial was 1 s. The inter-trial interval was 2 s. The number of days in which monkeys learned to accomplish this task with a criterion of

80% correct fixation (> 1 s) within a session was counted for statistical comparisons. Note that not all cynomolgus monkeys were able to accomplish the task at Stage 1.

Stage 2: Color discrimination. After the animal maintained a 500 ms fixation on a red circle on the screen, green and blue circles were simultaneously presented randomly on the left or right hand side of the touch screen. At the beginning of this stage, only one colored (green) target was rewarded. The monkey had to touch the correct color within 2 s to receive a 500ms reward (movie S5). The inter-trial interval was set at 1500 ms. Monkeys were trained for 2 weeks to reach stable performance, and then switched to another color (blue) reward session. Performance in each session was defined as the ratio of correct and total trials. Reaction time in correct trials was calculated for each monkey. The number of sessions taken to achieve an 80% correct rate within a single session was recorded. After acquiring the ability to differentiate green or blue colors for reward in different sessions, animals moved on to the next stage of training.

Stage 3: Reversal learning test. At the beginning of this stage (~2 weeks), monkeys took turns performing the green-reward session and blue-reward session on each experiment day to reinforce reversal learning and adapt to the intensity of training. Between sessions, there was no cue or clue provided to monkeys to indicate color change for juice reward. Monkeys had to discover this rule switch by trial and error. In the final test phase (~ 2 weeks), each monkey was tested with 10 sessions per day, each session containing 80 trials. The inter-session-interval was approximately 1 min. Within a session, one color (green or blue) was assigned for juice reward (movie S6-7). Between sessions, the color assigned for reward was randomly switched and counterbalanced among different monkeys. The percentage of correct trials in every ten trials performed after each switch was analyzed. Errors during discrimination could be more precisely differentiated as perseverative errors and regressive errors to further quantify the cognitive flexibility in switching session (*5*). Perseverative errors were responses in which monkeys continue to select the previously rewarded color following negative feedback but *prior* to the first correct selection after rule switch, suggesting a failure to switch to a new response. Regressive errors were responses in which monkeys continue to select the previously rewarded color *after* the first correct selection, thus representing an inability to maintain a new response. Attention level of the subject during this testing stage was examined by counting the omitted trials in each session, due to inability to overcome non-reward experience (*6*).

### Animal preparation

#### General anesthesia

For both EEG recording and fMRI scanning, animals were prepared and maintained in a stable brain state under light anesthesia. The animal preparation procedure was conducted in a manner similar to our previous work (*7, 8*). Induction of anesthesia was achieved by intramuscular injection with midazolam (0.25 mg/kg, Nhwa Pharma Co., Ltd., China) before EEG recording or with ketamine (10 mg/kg, Gutian Pharma Co., Ltd., China) before MRI scanning sessions, supplemented with atropine sulfate (0.05 mg/ kg, Shanghai Harvest Pharma Co., Ltd., China) to decrease bronchial and salivary secretions. After intubation, animals were ventilated with a mixture of isoflurane (2-2.5%, Lunan Pharma Co., Ltd., China) and oxygen via either a standard ventilator (CWE, Inc., Ardmore, PA, USA) outside the scanner room or an MRI-compatible ventilator (CWE Inc., Weston, Wisconsin) inside the scanner room. Macaques were maintained with intermittent positive-pressure ventilation to ensure a constant respiration rate (25-35 breaths/min). The concentration of isoflurane was adjusted based on continuously monitored vital signs, including blood oxygenation, electrocardiogram (ECG), rectal temperature (Small Animal Instruments, Inc., Stony Brook, New York), respiration rate and end-tidal $CO_2$ (Smiths Medical ASD Inc., Dublin, Ohio). Oxygen saturation was kept over 95% and body temperature was kept constant using a heated water blanket (Gaymar Industries Inc.,

Orchard Park, New York). Lactated Ringer's solution was given with a maximum rate of 10 ml/kg/hour during the anesthesia process (*9*).

**Anesthesia maintenance**

To monitor the brain state during the experiment, animals were further prepared with a MRI-compatible EEG recording system. To ensure data quality, we shaved the hair from the animal scalp and left back near the heart, and thoroughly cleaned the skin with abrasive gel and alcohol swabs to remove oil and dirt. A custom EEG cap made of stretchable materials with a two-panel design was fitted over the scalp. The junction between the anterior and posterior panels was aligned with the line between the ears, while the central electrodes were aligned to the midline of the head. The cap was then fastened by a chinstrap. An ECG electrode was attached close to the heart to facilitate off-line removal of cardio-ballistic artifacts. To increase the signal-to-noise ratio, we injected conductive gel and ensured a low impedance (<5 kilo-ohms) at each electrode. The electrode filling holes were subsequently covered with medical tape to prevent the gel from drying out during recording. After setting up the EEG cap, animals were restrained within the water blanket in a sphinx-like position with the head protruding and facing forward. Animal head was secured using a custom-built MRI-compatible stereotaxic frame after local anesthetic (5% lidocaine cream) was applied around the ears to block peripheral nerve stimulation.

In light of the anesthesiologist's instructions (SG, JC and YW), anesthesia was maintained using the lowest possible concentration of isoflurane gas during data acquisition. Isoflurane was selected for anesthesia maintenance as numerous studies have demonstrated that stable neural activity and functional connectivity patterns under a narrow range of medium level isoflurane (e.g., ±0.25%) are suitable for anesthetized nonhuman primate investigations (*7*, *10-12*). The concentration of isoflurane was adjusted based on both vital signs and EEG signatures.

Neither paradoxical excitation nor burst suppression, indicating minimally conscious and deep anesthesia states, respectively, was observed during data collection (*13*). Note that our intention was to equate the level of physiological anesthesia across animals and not the level of anesthetic gas concentration. Slight individual differences in physiology meant that slight differences in anesthetic gas concentrations were needed to impose a similar level of anesthesia on different monkeys (*14*). Within the range of isoflurane levels used in the current study, consistent patterns of functional coupling between distant brain areas have been reported in prior monkey fMRI studies (*12*, *15*) and demonstrated in our work as well (*7, 8*).

**Monkey EEG data acquisition**

Spontaneous neural activity was collected from five TG and sixteen WT monkeys in the EEG laboratory. EEG scalp recordings were acquired with the BrainVision Recorder software using a BrainAmp MR amplifier and a 28-channel EEG cap customized for macaques (Brain Products GmbH, Gilching, Germany) with sintered Ag/AgCl ring electrodes. The spatial distributions of electrodes over the brain were obtained in high-resolution T2-weighted MRI images using a three-dimensional turbo-spin-echo sequence (TR = 3000 ms; TE = 370 ms; field of view = 128 × 88 mm; acquisition voxel size = 0.5 × 0.5 × 0.5 $mm^3$; 30 sagittal slices), as demonstrated in Fig. 4A. Twenty-one active electrodes were projected from scalp to cortex (Fig. 4B) and labeled according to the closest cortical area (a complete list of anatomical labels in table S5). EEG signal was sampled at 5000 Hz with the resolution of 0.5 μV per bit and measuring range of ±16 mV. Due to a small number of TG subjects, repeated EEG measurement was performed such that 11 sessions were acquired from 5 TG monkeys. For each recording session, 6 to 10 runs lasting 7 min each were collected. Details of run numbers and physiological parameters for each animal are listed in table S4.

**Monkey MRI data acquisition**

MRI images of five TG and eleven WT monkeys were acquired at the Institute of Neuroscience on a 3T whole-body scanner (Trio; Siemens Healthcare, Erlangen, Germany) running with an enhanced gradient coil insert (AC88; 80 mT/m maximum gradient strength, 800 mT/m/s maximum slew rate). A custom-built 8-channel phased-array transceiver coil was used for animal imaging sessions. Whole-brain resting-state fMRI data were collected using a gradient-echo echo-planar sequence (TR = 2000 ms; TE = 29 ms; flip angle = 77°; slices = 32; matrix = 64 × 64; field of view = 96 × 96 mm; 1.5 × 1.5 $mm^2$ in plane resolution; slice thickness = 2.5 mm; GRAPPA factor = 2). For each session, 5 to 10 runs were acquired and each run consisted of 200 functional volumes. A pair of gradient echo images (echo time: 4.22 ms and 6.68 ms) with the same orientation and resolution as EPI images were acquired to generate a field map for distortion correction of EPI images. High-resolution T1-weighted anatomical images were acquired using a MPRAGE sequence (TR = 2500 ms; TE = 3.12 ms; inversion time = 1100 ms; flip angle = 9°; acquisition voxel size = 0.5 × 0.5 × 0.5 $mm^3$; 144 sagittal slices). Six whole-brain anatomical volumes were acquired and further averaged for better brain segmentation and 3D cortical reconstruction. Brain states were monitored by simultaneously recorded EEG signals, such that functional scans were collected under light to intermediate states of general anesthesia (*13*). In the present experimental design, we did not intentionally collect different numbers of runs from each subject. In practice, however, the actual scan time for each experiment varied with the physiological status of different animals on that day, as is the case in most animal fMRI studies (*12*, *15*, *16*). Furthermore, post hoc analysis in most imaging studies, including the present work, removes runs that show erratic vital signs, image artifacts or flawed small-worldness properties (for graph analysis of brain networks). These factors all led to different runs used in the final analysis, which included a total of 45 runs from TG and

99 runs from WT monkeys. Details of run numbers and physiological parameters for each animal are listed in table S8.

### Human MRI data

The human MRI data were obtained from the Autism Brain Imaging Data Exchange (ABIDE) (http://fcon_1000.projects.nitrc.org/indi/abide/). We screened human data based on two principles: (1) to make the data as homogenous as possible and (2) to make human data as well matched with the monkey data as possible. To reduce the notorious heterogeneity within human ASD, we conducted screening based on basic demographic and diagnostic information provided in the ABIDE database (*17*). The inclusion criteria for patients who were enrolled in the present study are briefly described as follows: (1) right-handedness, (2) age between 12~18 years, (3) a full-scale IQ (FIQ) score between 80 and 130, (4) a diagnosis of autism rather than Asperger syndrome or PDD-NOS according to the Diagnostic and Statistical Manual of Mental Disorders, Fourth Edition, Text Revision (DSM-IV-TR), (5) available site-matched typically developing controls (TDCs). The age limit was imposed so that the screened human matched with the monkey in terms of development stage. Finally, 90 adolescent individuals with autism (thereinafter, we use autism rather than ASD to refer to this patient cohort) and 140 TDCs matched for age, handedness, FIQ, and data source site were determined in order to enable an amenable comparison with the monkey neuroimaging data (*18*). Further details of the human sample are specified in table S11.

### EEG data analysis

### EEG preprocessing

EEG data analyses were conducted in MATLAB R2012a (The MathWorks). The preprocessing steps were implemented using custom-written scripts and routines from EEGLAB (*19*). Firstly, bad channels were identified via both visual inspection and automatic detection (using the probability and kurtosis criteria in EEGLAB with a standard deviation of 5) and excluded from further analysis. The data was then down-sampled to 1000 Hz (using the pop_resample function from EEGLAB). We chose a relatively high sampling rate to ensure good estimations of high-frequency activity in time-frequency analysis (*20*). The signal was subsequently high-pass filtered to remove slow drift with a Hamming-windowed zero phase finite impulse response filter at 1 Hz (using the pop_eegfiltnew function from EEGLAB). The filter order was optimized by the function and automatically set to 3300, with 6 dB attenuation at a cutoff frequency of 0.5 Hz. We used a sine-wave-fitting method provided in the CleanLine toolbox (*21*) to reduce electrical line noise at 50 Hz and its harmonics, in which noise estimation and removal was conducted separately for each channel. The amplitude and phase of a deterministic sinusoid of a specified frequency were estimated and statistically tested via Thomson's $F$-test (*22*). The exact line frequency was determined within 50 ± 2 Hz by maximizing $F$-statistics. If the amplitude of line noise frequency was significant ($P < 0.01$), the corresponding time-domain sinusoid was reconstructed and subtracted from the data (*23*). For the removal of pulse or cardio-ballistic artifacts, we employed the method of weighted average artifact subtraction (*24*), in which R-peak corresponding to each heartbeat was automatically detected and manually adjusted. Based on these peaks, an average artifact template for each channel was constructed over 21 consecutive heartbeat events, with less weighting on far away events (weighting factor = 0.9). The cardio-ballistic artifact was then removed by subtracting the artifact templates from the data. Visual inspection was meticulously conducted to ensure no residual artifacts due to incomplete artifact removal or other idiographic artifacts. After preprocessing, a total of 76 TG and 102 WT runs were included in the final analysis (table S4).

### Time-frequency analysis

**Time-frequency decomposition.** To estimate frequency domain features from resting state EEG, time-frequency representation of the preprocessed data was calculated for frequencies ranging from 1 to 100 Hz, using the ft_freqanalysis function from the Fieldtrip toolbox (*25*). Multi-taper transformation based on discrete prolate spheroidal sequences (*22*) was applied to windows of 4 s length that were moved over data in steps of 0.05 s. This method involves the multiplication of data segments with multiple tapers before the Fourier transform. Tapering effectively concentrates spectral estimates across a specified frequency band, with different tapers focusing on different parts of the resulting spectrum. We chose a spectral smoothing of ±1 Hz, which made the number of tapers *K* to be seven:

$$K = 2 * T * W - 1$$

where *T* is the segment length and *W* is the smoothing frequency.

Time-domain data were convolved with each taper and then Fourier transformed, giving a complex Fourier coefficient $\tilde{x}_k(f)$ for each frequency *f*:

$$\tilde{x}_k(f) = \sum_{t=1}^{N} w_k(t) x_t e^{-2\pi i f t}$$

where $x_t$ is the time series of the signal under consideration and $w_k(t)$ is the k-th taper function. The multi-taper estimates for the spectrum $s_x(f)$ and the cross-spectrum $C_{xy}(f)$ are given by:

$$s_x(f) = \frac{1}{K} \sum_{k=1}^{K} |\tilde{x}_k(f)|^2$$

$$C_{xy}(f) = \frac{1}{K}\sum_{k=1}^{K} \tilde{x}_k(f)\tilde{y}_k(f)^*$$

where $\tilde{x}_k(f)$ and $\tilde{y}_k(f)$ indicate Fourier coefficients from electrode sites x and y, and the asterisk indicates the complex conjugate of a complex value. For each EEG dataset, the spectrum at each recording site and cross-spectrum between each pair of sites were acquired for every time window, with a frequency resolution of 0.25 Hz.

**Spectral power analysis.** For power analysis, spectrograms representing temporal evolution of spectrum were segmented into 10 s non-overlapping epochs and averaged across epochs to obtain higher signal-to-noise ratio. Power spectra were calculated by taking median values of averaged spectrogram across time. To reduce inter-subject variability, the relative power was acquired by dividing the absolute power by the total power across the spectrum. The relative spectral power was categorized into six canonical frequency bands (*26*), namely delta (1–4 Hz), theta (4–8 Hz), alpha (8–12 Hz), beta (12–30 Hz), low gamma (30–60 Hz), and high gamma (60–100 Hz).

**Phase synchrony analysis.** For neural connectivity analysis between different recording sites, phase synchronization measures were estimated from a cross-spectrogram representing cross-spectrum at different times. An important challenge for accurate assessment of phase synchronization and relevant EEG-based neural connectivity is the issue of volume conduction (*27*), which refers to the fact that the electrical potentials generated by neuronal activity are not only measured in the direct vicinity of neuronal sources but can also be measured at distant sites. It is therefore suggested to confine the analysis to non-instantaneous correlations, such as the phase-lagged part of coherence (*27*). In the present study, we analyzed phase-lagged synchronization using de-biased weighted phase-lag index (dwPLI) (*28*). For two EEG signals at electrode sites *x* and *y*, phase spectrum $\phi_{xy}$ at frequency *f* can be estimated from cross-spectrum $C_{xy}(f)$:

$$\phi_{xy}(f) = \arctan(-\frac{\Im\{C_{xy}(f)\}}{\Re\{C_{xy}(f)\}})$$

where $\Im\{.\}$ and $\Re\{.\}$ indicate the imaginary and real part of a complex value. A non-zero imaginary component thus indicates a non-zero phase difference ('lag') between two signals. As the time window moves over the data, a series of phase-lag values can be estimated. The distribution of these phase-lag values is expected to be symmetric when it is flat (no coupling) or centers around 0 mod $\pi$ (influence of common source or active reference). In contrast, asymmetry of the distribution implies the presence of a consistent, nonzero phase-lag between the signals. The asymmetry level could be estimated via phase-lag index (PLI), which is defined as the absolute average value of the sign of the imaginary part of $C_{xy}(f)$ (hereafter referred to as $C$):

$$PLI = |E\{sgn(\Im\{C\})\}|$$

where $E\{.\}$ is the expected value operator. PLI ranges from 0 to 1, with 0 indicating no phase-lagged coupling and 1 indicating constant phase difference (*29*). The weighted PLI (wPLI), which is less sensitive to noise than PLI (*28*), extends the original PLI in that it weights the contribution of observed phase differences by the magnitude of the imaginary component of the cross-spectrum:

$$wPLI = \frac{|E\{\Im\{C\}\}|}{E\{|\Im\{C\}|\}} = \frac{|E\{|\Im\{C\}|sgn(\Im\{C\})\}|}{E\{|\Im\{C\}|\}}$$

Both weighted and original PLI are susceptible to sample-size bias, i.e. the estimation increases as sample size (in this case, the number of time windows) decreases. The unbiased version of PLI (ubPLI) can be calculated as the average of all pairwise products of signs

$$ubPLI = \mathrm{E}\{\mathrm{sgn}\left(\Im\{C_j\}\right) * \mathrm{sgn}(\Im\{C_k\})\}$$

where indexes $j$ and $k$ indicate different time windows. Similarly, the de-biased (i.e., some sample-size bias remains) version of wPLI (dwPLI) can be calculated as:

$$dwPLI = \frac{\sum_{j=1}^{N} \sum_{k \neq j} \Im\{C_j\}\Im\{C_k\}}{\sum_{j=1}^{N} \sum_{k \neq j} |\Im\{C_j\}\Im\{C_k\}|}$$

It has been demonstrated that dwPLI is more robust against volume conduction, noise and sample-size bias than other neural connectivity measures (*28*). For each EEG dataset, a 21 × 21 matrix of dwPLI values was generated for each frequency bin. Band-specific synchronization strength was calculated by averaging dwPLI values across frequency bins within the band.

**EEG statistical analysis**

After a linear regression model with age, age$^2$, and gender as covariates was applied to estimate age and gender effects, group comparisons of EEG power and synchronization strength were conducted separately for individual frequency bands. Differences in relative power between the two groups were tested using independent two-sample *t*-tests, with Bonferroni correction for multiple comparisons. Effect size, an estimate of population parameters that are independent of sample size and other design decisions, was estimated for each channel via *Hedges' g* value. Effect sizes were divided into three levels: small, medium, and large, each corresponding to a *Hedges' g* value greater than or equal to 0·2, 0·5, and 0·8. Effect size (*Hedges' g* value) was estimated in the Measures of Effect Size (MES) Toolbox (*30*).

As for the between-group differences in neural synchronization, the covariates-free dwPLI matrices were first logarithm transformed to normalize the highly skewed distribution and then compared using the network-based statistic approach (*31*). This approach copes with the multiple comparison problems of comparing connectivity data with $N$ nodes and $N \times (N-1)/2$ connections (or edges). The correction strategy is analogous to cluster-based strategies used in voxel-wise MRI data analysis by evaluating the null hypothesis at the level of connected subnetworks rather than individual connections. In this method, subnetworks were identified with a strategy similar to identifying significant clusters in fMRI analysis (*31*). Specifically, an independent two-sample $t$-test was applied to each connection in the covariates-free dwPLI matrix. A primary threshold ($P$ < 0.005) was applied to the connection-level statistical map. The remaining connections may form connected components (with the number of connections > 3). A non-parametric permutation approach (10,000 times) was then used to ascribe a $P$-value to each connected component based on its size. For each permutation, the group labels were randomly exchanged, and $t$-statistics were re-calculated. Subsequently, the same threshold was applied to define supra-threshold connections, after which the maximal component size was calculated. Finally, the $P$-value of an observed component of size $Q$ was estimated as the percentage of the permutations that had a higher maximal component size larger than Q. All connections within components with a significant group difference ($P$ < 0.05) were considered significant. Effect sizes (*Hedges' g* value) were estimated for all significant connections.

The overall abnormality in neural synchronization of an individual TG monkey relative to the averaged WT control was estimated via Manhattan distance based on identified connections with significant group difference, which reflects the sum of the absolute differences of disrupted neural circuits (*32, 33*). The formula for Manhattan distance is $\sum_{i=1}^{n} |x_i - y_i|$, where $x$ and $y$ are two vectors and $i$ indexes the $n$ elements of the vectors. The vectors here were all connections within significant components. To evaluate the significance of the calculated Manhattan distance, 5 null distributions corresponding to 5 TG monkeys were generated via 10,000 shuffles, during which labels (TG or WT) were randomly shuffled to produce two random groups; Manhattan distances between each of the 5 randomly-labeled TG monkeys and an averaged WT monkey were then calculated. The number of permutations where random

Manhattan distance is not smaller than in the real condition was divided by 10,000 to obtain a $P$ value for each monkey. We consider $P < 0.05$ to be statistically significant.

Associations between gene (*MECP2* copy number), circuit (Manhattan distance based on beta synchronization strength of all abnormal connections), and behavior (general locomotion time and repetitive index in home cage, locomotion change induced by familiar peer separation, and error rates in reversal learning task) through calculating their Spearman's correlation coefficients in TG monkeys.

**MRI data analysis**

**Network construction of monkey and human brains**

Functional images of monkey and human brains were preprocessed using exactly the same strategy, which included slice timing correction, motion correction, coregistration with individual T1-weighted image, normalization to corresponding standard space, reslicing and spatial smoothing, regression of nuisance signals, removal of linear drift and temporal filtering (0.01 - 0.1 Hz). Specifically, the preprocessing of the monkey data were done using the SPM 8.0 toolbox (http://www.fil.ion.ucl.ac.uk/spm) and the FMRIB Software Library toolbox (FSL; http://www.fmrib.ox.ac.uk). The first 10 volumes were discarded. The field map images of each participant were then applied to compensate for the geometric distortion of EPI images caused by magnetic field inhomogeneity using FSL FUGUE. After slice timing correction and motion correction, the corrected images were normalized to standard space of the monkey F99 atlas (http://sumsdb.wustl.edu/sums/macaque more.do) using an optimum 12-parameter affine transformation and nonlinear deformations, and then resampled to 2-mm cubic voxels and spatially smoothed with a 4 mm full-width at half-maximum (FWHM) isotropic Gaussian kernel. Six head motion parameters, ventricle and white matter signals were removed from the smoothed volumes using linear regression. Linear drift of the volumes was removed and a temporal filter was performed. On the other hand, the preprocessing of human data was performed by the Preprocessed Connectomes Project (PCP, http://preprocessed-connectomes-project.org/abide/index.html) using the Data Processing Assistant for Resting-State fMRI (DPARSF) Toolbox. Preprocessing steps included slice timing correction, motion correction, spatial normalization into MNI space, reslicing to 3 × 3 × 3 mm voxels and smoothing with a Gaussian kernel (FWHM = 6 mm). Friston-24 parameters of head motion, white matter and ventricle signals were regressed out, followed by linear drift correction and temporal filtering. For more details, readers may refer to the description in the PCP (http://preprocessed-connectomes-project.org/abide/dparsf.html).

The cortical organization of both monkeys and humans was parcellated according to the Regional Map template (*34, 35*). As Regional Map parcellation does not include subcortical regions, subcortical parcellation for the two species was added on the basis of the INIA19 (*36*) and Freesurfer templates (*37*), respectively. This generated a whole brain template with a total of 94 regions of interest (ROIs) for both monkeys and humans (see table S7 for a complete list of all anatomical labels). Pearson's correlation coefficients between the mean time courses of any pair of regions were calculated to represent their functional connectivity, resulting in a 94 × 94 connectivity network matrix. Fisher's Z-transformation was then applied to the connectivity matrix which was subject to a covariates regression procedure before subsequent analysis. For the monkey data, age, $age^2$ and gender were regressed out of connectivity matrices using a general linear model. For the human data, covariates including age, $age^2$, gender, FIQ and scanning sites were regressed out of connectivity matrices using a general linear model.

**Parsing heterogeneity in clinical cohorts**

It is well known that substantial heterogeneity exists in the etiology and phenotypes of ASD. We used data-driven stratification of functional connectivity to reveal clusters or subgroups present across autism and TDCs. Emerging evidence has demonstrated that healthy and ill individuals may share biological commonalities and form subgroups regardless of their clinical status (*38*), using either behavior (*39*, *40*) or imaging measures such as functional brain networks (*41-43*). A total of 230 human samples (90 autism and 140 TDCs) were used to generate a 230 × 230 similarity (spatial correlation) matrix by calculating Pearson's correlation coefficient between the brain networks (lower triangle of network matrix) of any two participants. This inter-individual similarity matrix was rarefied as the maximum threshold was determined (Pearson's $r$ = 0.55, sparsity = 0.42) to keep the whole matrix connected. A community detection method was then applied, similar to that utilized in our recent work (*44*), to stratify human participants based on the inter-individual similarity of brain connectivity networks. This procedure was implemented using the Brain Connectivity Toolbox (*45*). Various types of algorithms have been developed to identify the optimal modular decomposition (*46*). These algorithms attempt to maximize modularity, $Q$, which reflects the difference in the degree of intra-modular edges between the observed graph and a random graph (*47*). The function of $Q$ can be written as:

$$Q(G) = \frac{1}{2m}\sum_{i \neq j}\left(A_{ij} - P_{ij}\right)\delta(M_i, M_j)$$

where $m$ is the total weight of edges in graph $G$; $A_{ij}$ is the weighted edges between $i$ and $j$; $\delta(M_i, M_j)$ is 1 if $i$ and $j$ are in the same module and 0 otherwise; $P_{ij}$ represents the probability that there would be an edge between $i$ and $j$ in a random rewired graph of $G$; $P_{ij} = k_i k_j / 2m$, where $k_i$ is node $i$'s degree. Newman's spectral optimization algorithm was used to decompose the similarity matrix. Hence, two communities or modules were detected automatically ($Q$ = 0.13), that is, these 230 participants were clustered into two subgroups. Detailed information about these two subgroups of human data are listed in tables S12 and S13.

After stratification, the within-cluster homogeneity between participants with autism and TDCs was statistically compared. The network similarities of one participant with all other participants from the same subgroup were averaged to get a mean value which was used to indicate the homogeneity of each subject with all other participants in the same experimental group. A permutation test was then applied to assess statistical differences in the homogeneity between subgroups by randomly shuffling all participants' labels (autism or TDC). The shuffling procedure was repeated 10,000 times, giving rise to 10,000 random differences that constructed a null distribution. A one-sided *P* value – the percentage of the permutations that had a larger (right side) or smaller (left side) group difference – was obtained. Using the same permutation procedure, homogeneity comparison between the entire cohort and two subgroups/clusters were also conducted.

**MRI statistical analysis**

Two-sample *t*-tests with two tails were applied to evaluate statistical significance of group differences. Edge-wise threshold of the significance level was set at $P = 0.001$, and cluster-level correction of $P < 0.05$ was applied to adjust the multiple comparison using the same network-based statistic as described in EEG statistical analysis (*31*). We continued to evaluate the extent of observed group effect by using effect size, measured by *Hedges' g* value, in addition to the common *P* value measure of statistical significance. The effect size of each brain node (*ESn*) was calculated by summing up the ESs of all edges with significant group difference connecting to this region. The *ESn* of brain regions were divided by the absolute maximum *ESn*, leading to a scale between -1 and 1. For the human data, effect size calculation and statistical tests were conducted on covariate-free brain networks for each subgroup, respectively. The overall measure of network dysfunction was evaluated via Manhattan distance and statistically tested via non-parametric permutation in a similar way as in EEG statistical analysis.

To assess the distribution of these disrupted edges within and between lobes, we used the standardized residuals (*48*) to adjust the bias caused by the Regional Map parcellation in which the number of nodes is not equal in different lobes. The standardized residuals (*Z* scores) are defined as the raw residuals (or the difference between the observed distribution and expected distribution) divided by the square root of the expected distribution which hypothesizes that all the edges are randomly distributed across lobes. We then used a nonparametric statistical test to estimate the significance of observed distribution. Specifically, the disrupted edges were randomly assigned across the whole brain network and the standardized residuals were then re-calculated. This procedure was repeated 5,000 times and the null distribution was generated. The percentage of the assignments that had a larger or equal *Z* score was defined as the *P* value. Bonferroni correction was applied for the correction of multiple comparisons.

To dissect the circuit of different phenotypic dimensions confined in disrupted neural circuits, we further explored functional connections correlated with each behavioral measure by calculating Spearman's correlation coefficient between original functional connectivity and behavioral metrics of all monkeys that were available. Edge-wise threshold of the significance level was set at $P = 0.01$ for spontaneous locomotion in home cage (data available for 16 monkeys) and $P = 0.05$ for behavioral measures from peer separation test and reversal learning task (data available for 8 and 9 monkeys, respectively). Cluster-level correction of $P < 0.05$ was applied to adjust for multiple comparisons (*44*). Particularly, edges with a Spearman's correlation *P* value lower than the threshold were defined as suprathreshold connections and decomposed into several connected components. The significance of each clustered component was estimated in a similar way as in network-based statistic via nonparametric permutation (5,000 permutations). For each permutation, phenotype performance was randomly shuffled, and the Spearman's correlation coefficient was recalculated for each edge. The same threshold ($P = 0.05$) was then applied to define the suprathreshold connections, and maximal component size in the set of suprathreshold links was recorded. The statistical significance of a connected component with size $S$ was computed as the percentage of the null distribution that had a maximal component size larger than $S$.

Manhattan distance between each TG and the averaged WT control was calculated based on the functional connectivity of edges within each subnetwork that was correlated with a specific behavior. Gene-circuit-behavior associations between *MECP2* copy number, circuit abnormality of each subnetwork (Manhattan distances), and the corresponding behavioral measure were assessed via Spearman's correlation. Meanwhile, the Manhattan distance between connectivity fingerprint of each autism case and averaged TDC was calculated to quantitatively assess the circuit abnormality in individual autism. The relationship between this Manhattan distance (representing overall abnormalities of connectivity fingerprint) and clinical scores of Autism Diagnostic Observation Schedule (ADOS), including the total score and sub-scores of communication, social interaction and stereotypic behavior, were evaluated using both Spearman's and Pearson's correlation coefficients. This correlation analysis was not performed for subgroup 2 because no group differences in functional connections survived the correction for multiple comparisons. Note that clinical scores of ADOS were only available for 29 and 25 participants from subgroup 1 and subgroup 2, respectively (tables S12 and S13).

**Cross-species comparison of connectivity fingerprints**

To seek potential pathological correspondence between abnormalities of neural circuits identified in *MECP2* duplication monkeys and patients with autism, we conducted cross-species similarity analyses by comparison of altered functional homologs at multiple spatial scales: the whole brain level (the entire ES matrices of monkeys and humans) (*49*), the brain lobe level (the entire ES matrices were categorized into eight key lobes), and the brain node level (the entire ES matrices were split into 94 brain regions). As the connectivity matrices, including the ES matrices, were symmetric, the calculation of Pearson's correlation coefficient between two species was based on the lower triangle of the matrix. The statistical significance threshold was set at $P < 0.05$ after Bonferroni correction for multiple comparisons.

20. M. X. Cohen, *Analyzing Neural Time Series Data: Theory and Practice*. J. Grafman,

Ed., Issues in Clinical and Cognitive Neuropsychology (The MIT Press, Cambridge, MA, London, UK, 2014).

29. C. J. Stam, G. Nolte, A. Daffertshofer, Phase lag index: Assessment of functional connectivity from multi channel EEG and MEG with diminished bias from common

sources. *Hum. Brain Mapp.* **28**, 1178-1193 (2007).

48. D. J. Sheskin, *Handbook of Parametric and Nonparametric Statistical Procedures: Third Edition*. (CRC Press, 2003).

49. M. W. Cole, D. S. Bassett, J. D. Power, T. S. Braver, S. E. Petersen, Intrinsic and task-evoked network architectures of the human brain. *Neuron* **83**, 238-251 (2014).

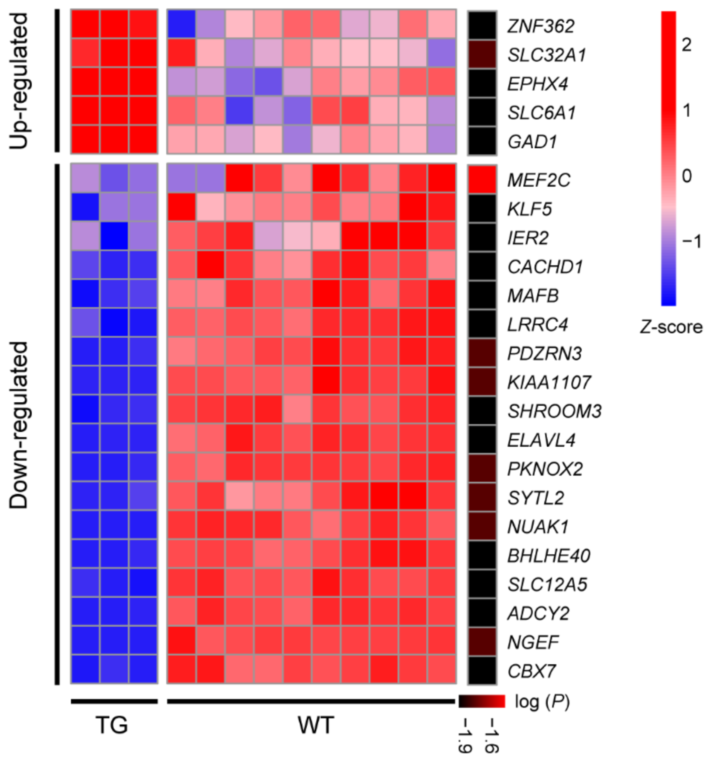


**Fig. S1. Heatmap for the expression of *MECP2* top correlated genes in the cortical regions of *MECP2* transgenic monkeys.** Significant level (log (*P*)) is indicated in the sidebar. Blue: low expression; Red: high expression.

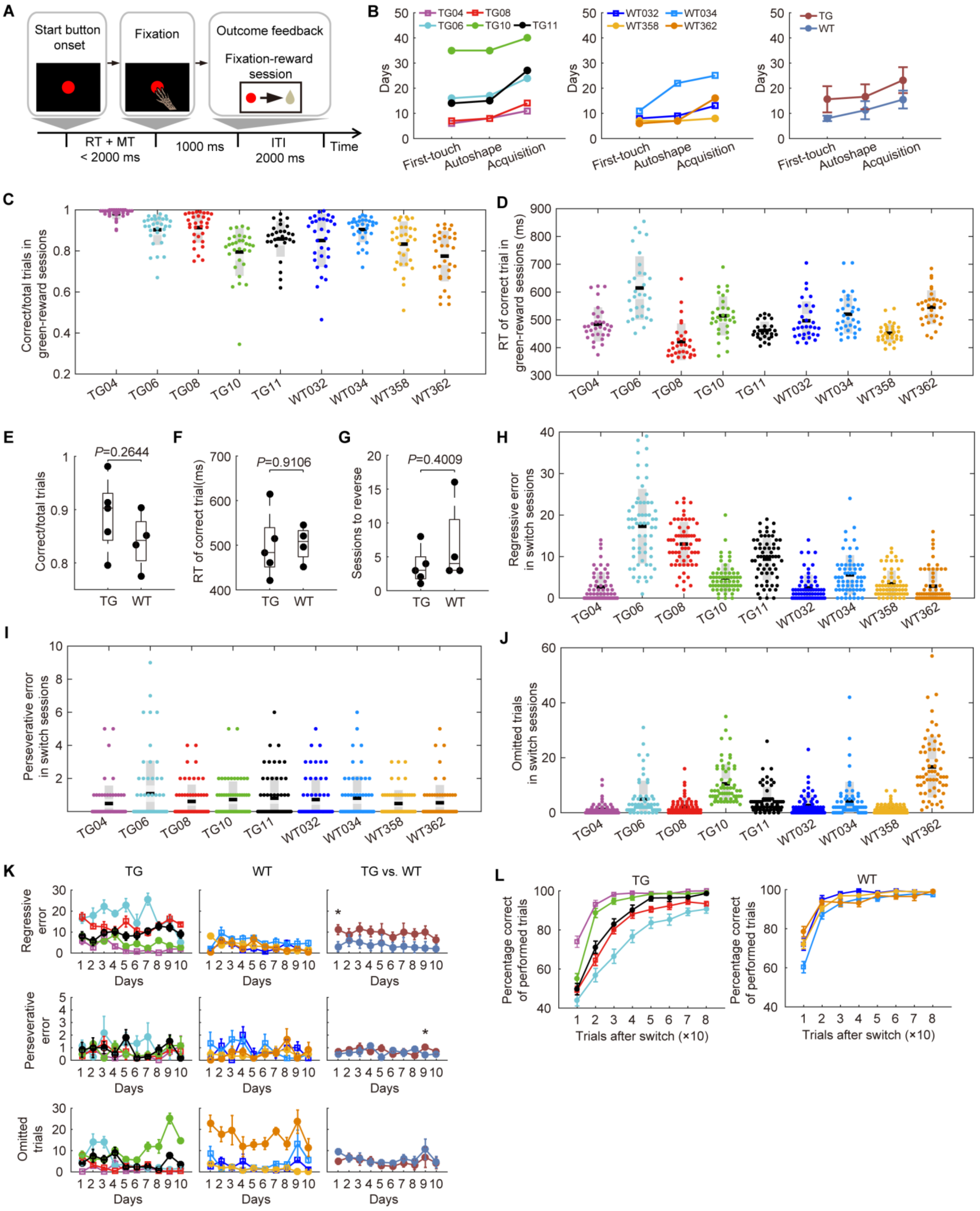
A
Start button onset
Fixation
Outcome feedback
Fixation-reward session
RT + MT < 2000 ms
1000 ms
ITI 2000 ms
Time
B
TG04 TG08 TG06 TG10 TG11
WT032 WT034 WT358 WT362
TG WT
Days
First-touch
Autoshape
Acquisition
C
Correct/total trials in green-reward sessions
D
RT of correct trial in green-reward sessions (ms)
TG04 TG06 TG08 TG10 TG11 WT032 WT034 WT358 WT362
E
P=0.2644
Correct/total trials
F
P=0.9106
RT of correct trial(ms)
G
P=0.4009
Sessions to reverse
TG WT
H
Regressive error in switch sessions
I
Perseverative error in switch sessions
J
Omitted trials in switch sessions
K
TG
WT
TG vs. WT
Regressive error
Perseverative error
Omitted trials
Days
L
TG
WT
Percentage correct of performed trials
Trials after switch (×10)

**Fig. S2. Behavior results of color discrimination and reversal learning task. (A)** Temporal sequence of events in stimulus-directed touching. After the start button appeared on the screen, subjects had to touch the button within 2000 ms to receive a reward. Subjects were rewarded for no longer than 1000 ms, as long as they fixed on the screen. The inter-trial interval was 2000 ms. RT, reaction time; MT, movement time; ITI, inter-trial interval. (**B)** Number of days until first-touch, autoshape and acquisition steps in stimulus-directed touching for five TG and four WT monkeys. **(C, D)** Scatter plot of individual performance **(C)** and reaction time in correct trials **(D)** in green-reward sessions. Each point denotes performance in each session. Black lines denote means. Dark gray and light gray boxes represent standard error of mean and standard deviation. **(E-G)** Group comparison of performance (**E**), reaction time of correct trials (**F**), and sessions to reverse (**G**) between TG and WT monkeys ($n$ = 5, TG; $n$ = 4, WT; two-tailed Student's $t$-test). (**H-J)** Scatter plot of regressive errors (**H**), perseverative errors (**I**) and omitted trials (**J**) in switch sessions. Each point denotes performance in each session. (**K)** Individual and group comparison of regressive errors, perseverative errors and omitted trials during 10 training days (*$P$ = 0.030 for comparison of regressive errors on the first day, *$P$ = 0.047 for comparison of perseverative errors on the 9$^{th}$ day, two-tailed Student's $t$-test). **(L)** Individual performance of percentage correct of performed trials. Error bars denote standard error of mean.

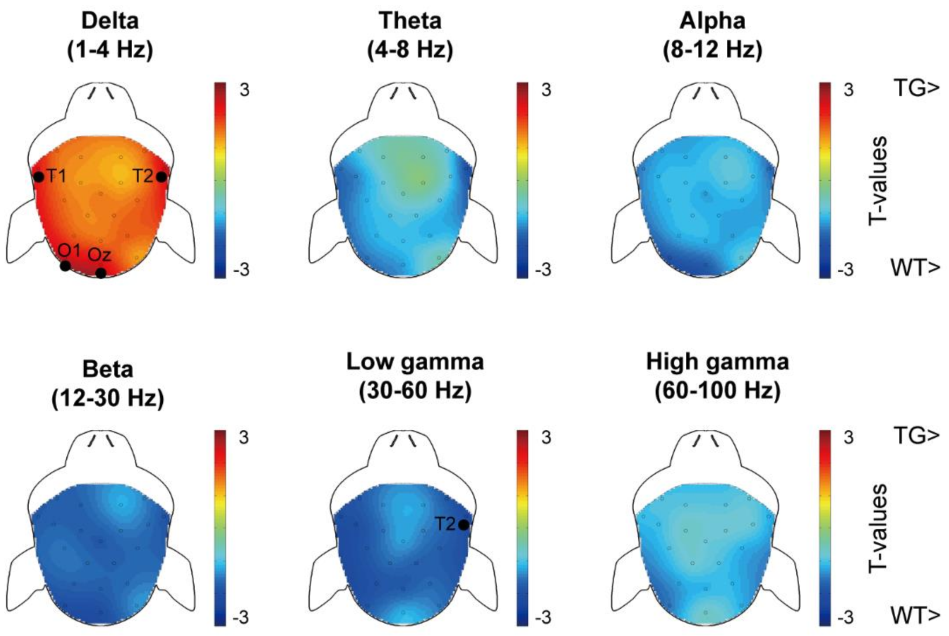


**Fig. S3. Topography of relative power differences between TG and WT monkeys for six frequency bands.** Effects of age and gender were regressed out before group comparison. The color scale stands for the *t*-statistic values. Black dots indicate the locations where group difference in relative power was observed at the significance level of $P < 0.05$ (uncorrected).

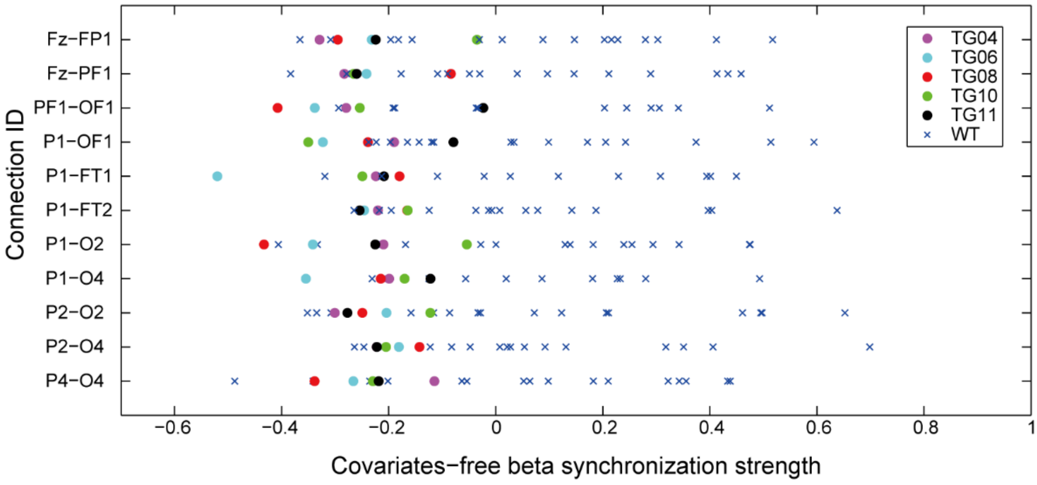


**Fig. S4. Strength of connections with significantly decreased beta synchronization in TG monkeys.** Solid circles in different colors represent individual TG monkeys ($n$ = 5). Blue cross markers represent individual WT monkeys ($n$ = 16). Full names of connection ID are listed in table S5.

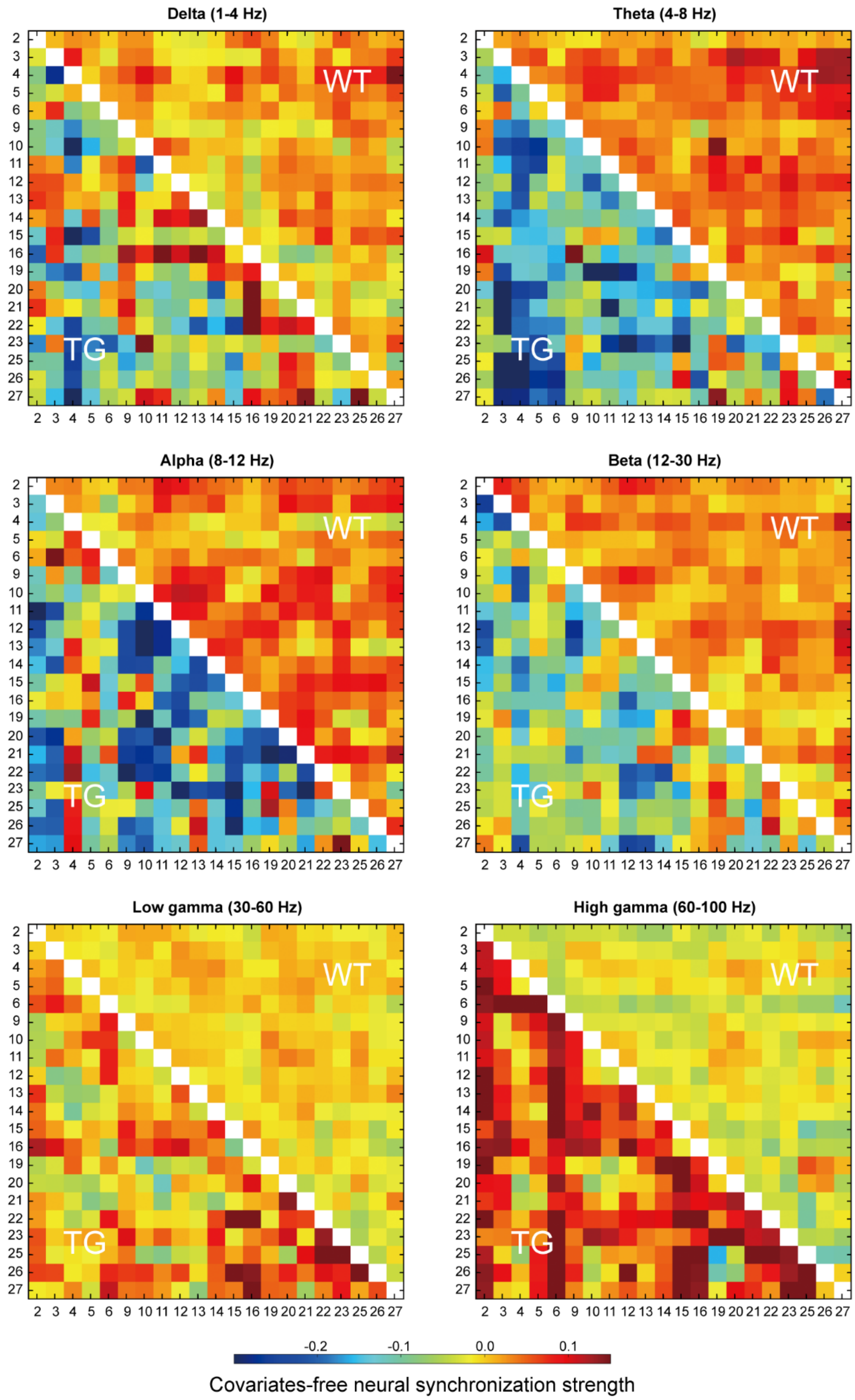


**Fig. S5. Averaged neural synchronization matrices for both TG and WT monkeys for six frequency bands.** The numbers indicate the electrode ID as in table S5. Color bar indicates dwPLI values.

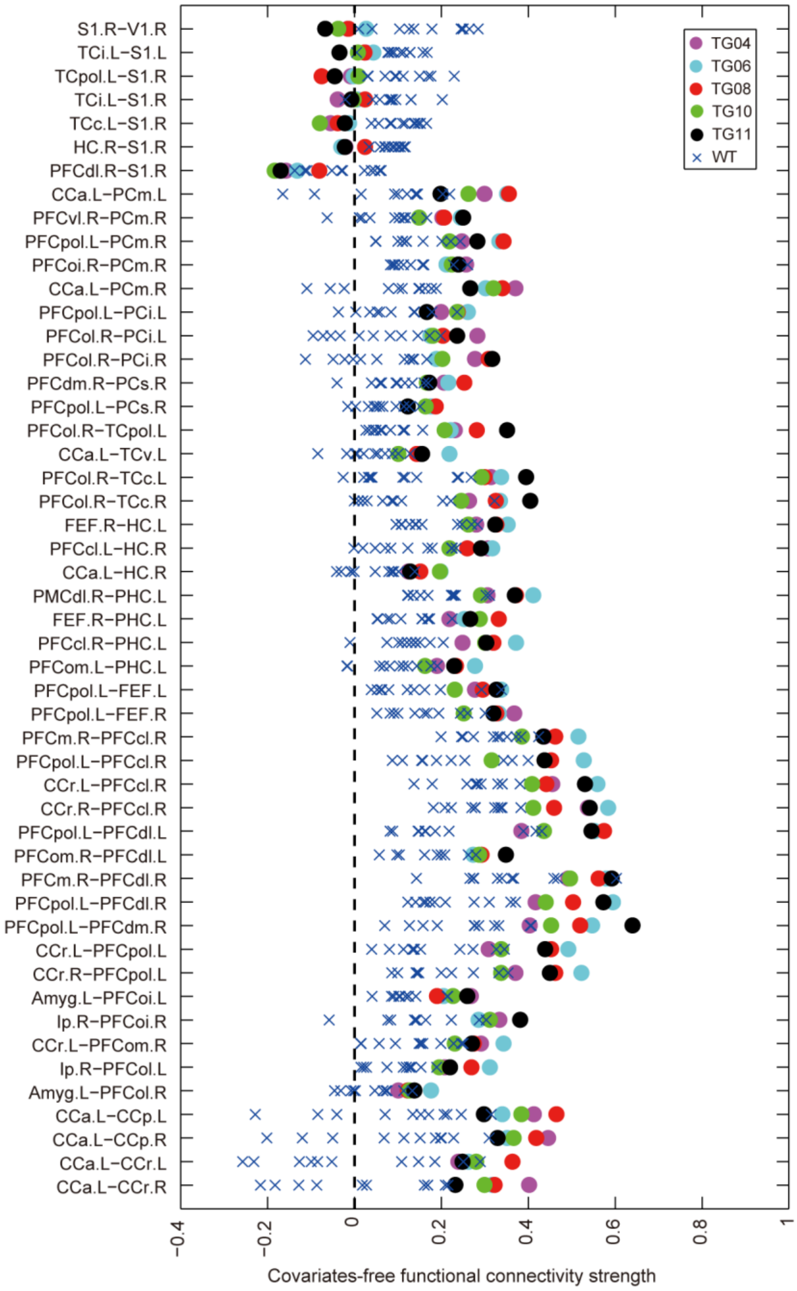


**Fig. S6. Scatter plot of abnormal connections in transgenic monkeys.** X axis indicates the magnitude of covariate-free functional connectivity. Solid circles in different colors represent individual TG monkeys ($n = 5$). Blue cross markers represent individual WT monkeys ($n = 11$).

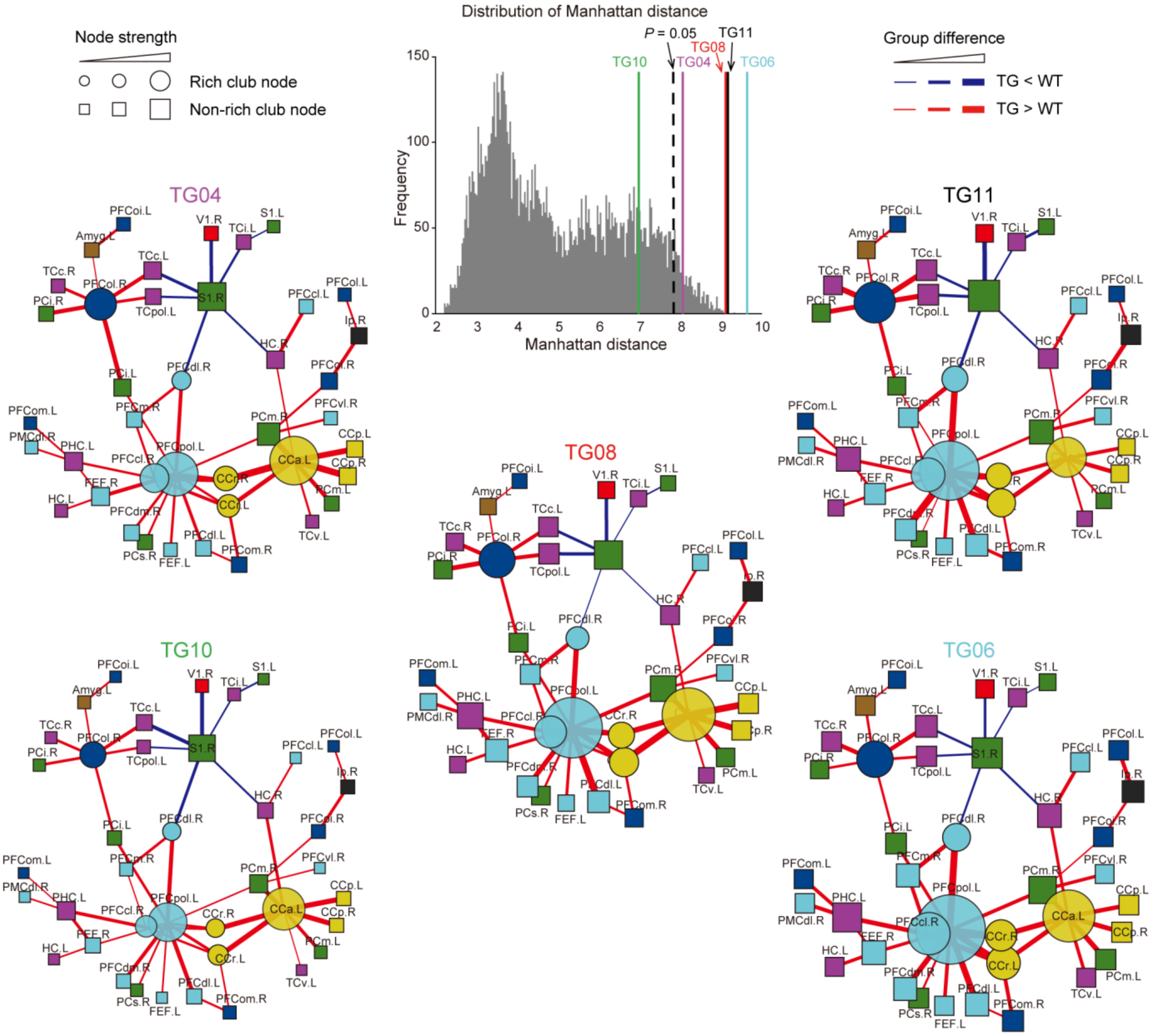

**Fig. S7. Connectivity fingerprint of individual transgenic monkeys.** The connectivity fingerprint of each TG monkey quantitatively assessed by Manhattan distance of disrupted neural circuits between individual TG and WT controls. The null distribution (gray) was generated by 10,000 permutations, with TG and WT labels randomly permuted in each round. All TG except for TG10 were evidently different from WT monkeys. The spider plots exhibit the connectivity fingerprint of each TG, in which the rounded nodes represent rich-club members. The color and thickness of the line between nodes indicate the sign (red: increase; blue: decrease) and the extent of difference in connectivity network of TG relative to WT monkeys. Node size indicates nodal strength, that is, the summation of weighted lines connecting to it.

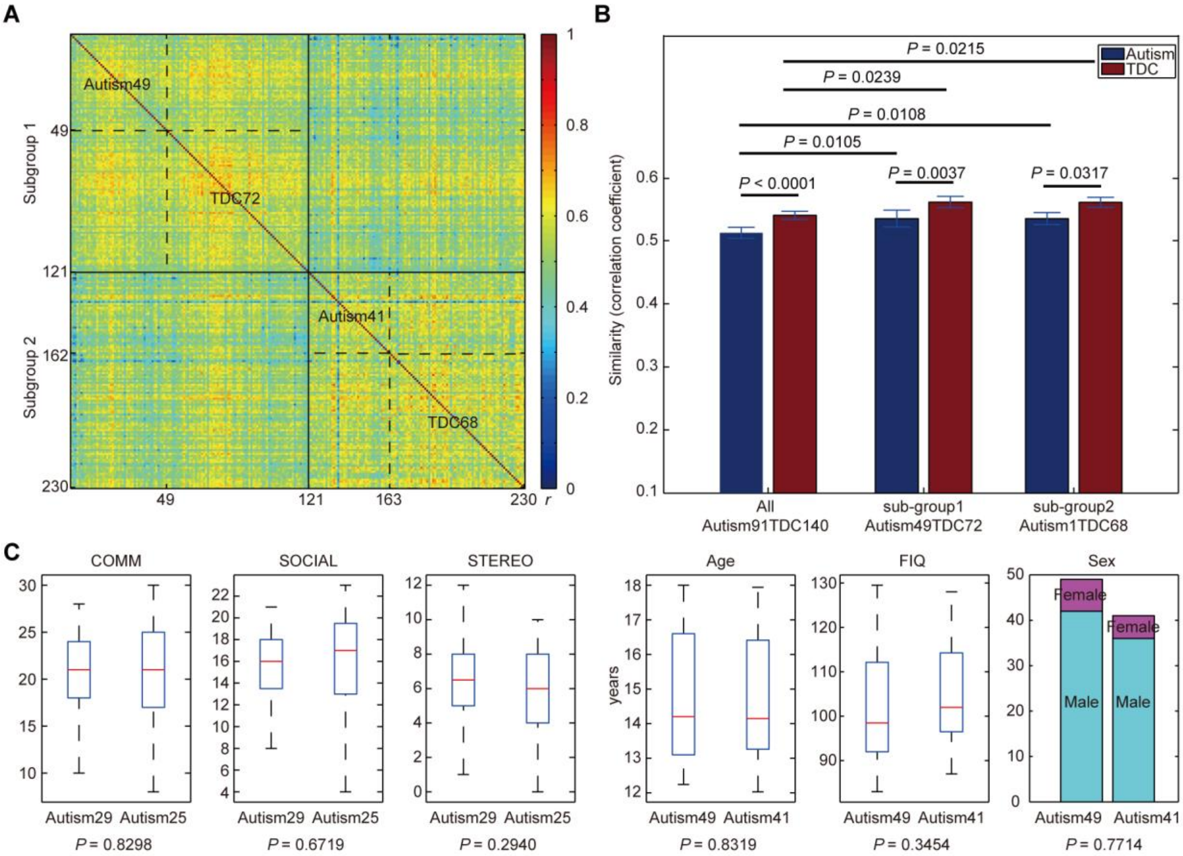


**Fig. S8. Parsing heterogeneity in clinical cohorts. (A)** An inter-individual network similarity matrix of 230 participants clustered by a community detection algorithm. Two sub-groups or clusters were derived out of this cohort, one with 49 individuals with autism and 72 TDCs (subgroup 1) and another with 41 individuals with autism and 68 TDCs (subgroup 2). Each row represents the similarities between the brain network of a single subject and that of all other subjects. Color bar, Pearson's correlation coefficients (*R*). **(B)** Group comparison of inter-individual network similarity. Brain network similarities of TDCs are always significantly higher than those of autism for both the entire cohort and two subgroups (10,000 times of permutation test). Brain network similarities of subgroups are significantly higher than those of autism and TDC groups, respectively (10,000 times of permutation test). Error bars, standard deviation. **(C)** Comparisons of demographic information and clinical scores of human autism between subgroup 1 and subgroup 2. Note that clinical scores are only available for some patients in both subgroups (29 and 25 for subgroup 1 and 2, respectively). In the boxplot, the central mark is the median and the edges of the box are the 25th and 75th percentiles. The whiskers extend to the most extreme data points. COMM, communication total sub-score of the classic ADOS; SOCIAL, social total score of the classic ADOS; STEREO, stereotypic behaviors and restricted interests total sub-score of the classic ADOS. FIQ, full-scale IQ.

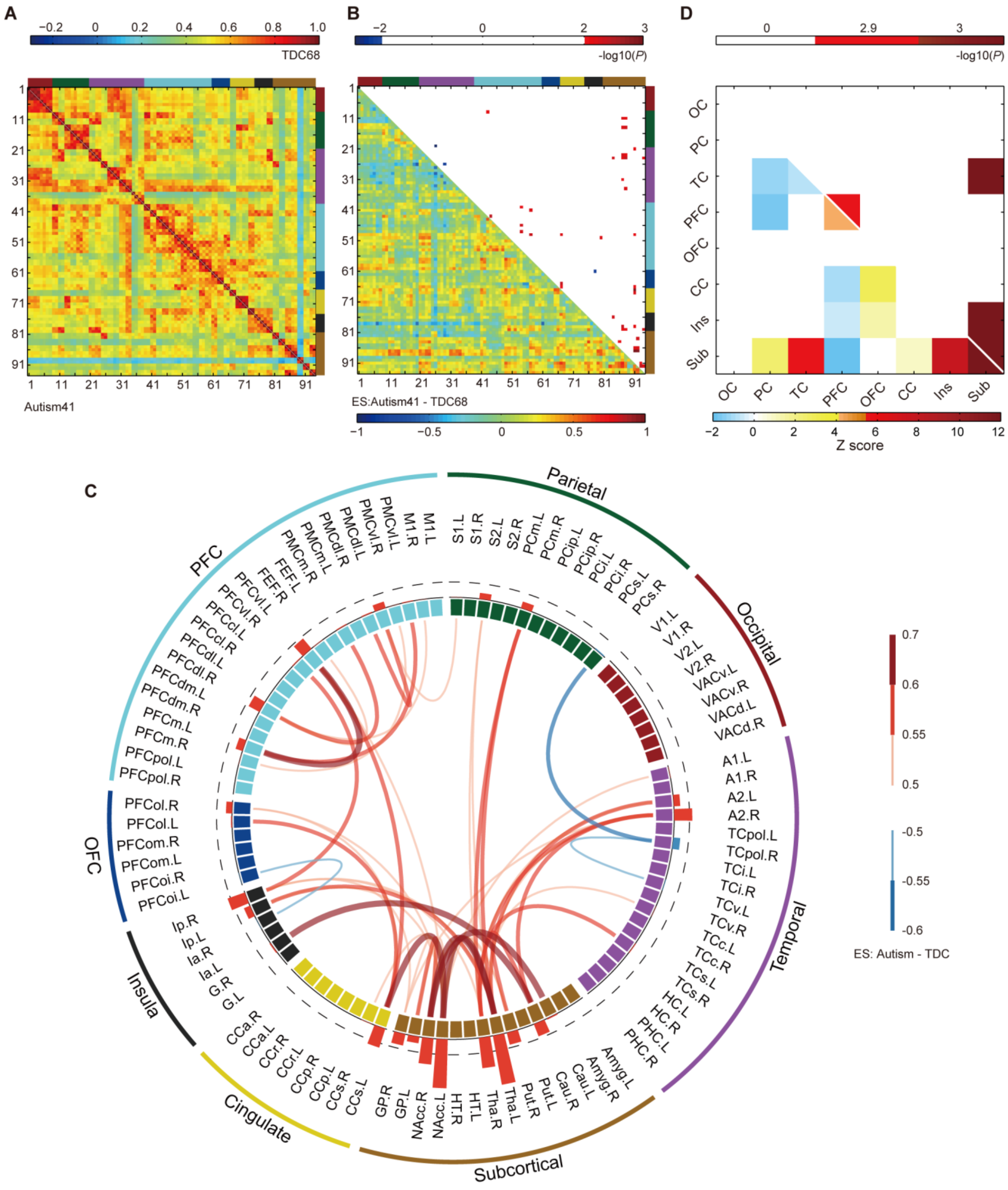

**Fig. S9. Abnormal functional connections of human autism in subgroup 2. (A)** Averaged functional connectivity networks for autism (left bottom) and TDC (right top). **(B)** Effect sizes

(ES) of autism versus TDC (left bottom) and statistically significant *P* values (right top, $P <$ 0.01, uncorrected). Note that the threshold of statistical tests is more liberal than that of subgroup 1 as no connections survived the correction for multiple comparisons. The colors on the top and right of the matrix in (**A**) and (**B**) indicate lobes that corresponding nodes belong to. **(C)** Disrupted functional connections and corresponding brain regions in autism. The red and blue bars in the interlayer indicate positive and negative *ESn*, respectively. For abbreviation and topology of brain nodes refer to table S7. **(D)** Distribution of disrupted connections within and between lobes (left bottom) and statistical significances (right top, $P < 0.05$, Bonferroni correction). Distribution of disrupted connections was measured via standardized residuals (*Z* scores), similarly calculated in Fig. 5D. OC, occipital cortex; PC, parietal cortex; TC, temporal cortex; PFC, prefrontal cortex; OFC, orbitofrontal cortex; CC, cingulate cortex; Ins, insula; Sub, subcortical areas.

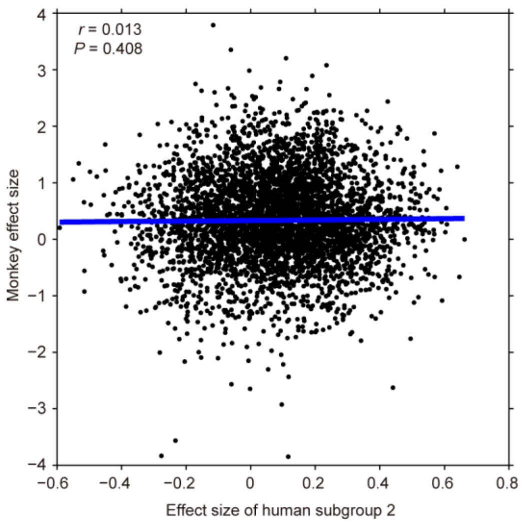


**Fig. S10. Spatial correlation between entire effect size (ES) matrices of TG versus WT monkeys and human autism versus TDC in subgroup 2.**

**Table S1. *MECP2* copy number and locomotion of 5 TG and 16 WT monkeys in home cage**

| ID | *MECP2* copy number | Sex | Age (years) | Repetitive circular locomotion (1,000s) | General locomotion (1,000s) | Repetitive index in locomotion [a] |
|---|---|---|---|---|---|---|
| TG04 | 1.0 | M | 4.7 | 0.97 | 2.54 | 0.38 |
| TG06 | 7.3 | F | 4.7 | 1.06 | 2.61 | 0.41 |
| TG08 | 2.9 | M | 4.7 | 1.49 | 2.42 | 0.62 |
| TG10 | 1.1 | F | 4.7 | 1.77 | 3.51 | 0.50 |
| TG11 | 1.9 | F | 4.5 | 0.06 | 2.82 | 0.02 |
| WT019 | - | M | 7.3 | 0.05 | 0.13 | 0.35 |
| WT026 | - | M | 7.1 | 0 | 0.49 | 0 |
| WT027 | - | M | 7.1 | 0.11 | 0.48 | 0.22 |
| WT030 | - | F | 4.7 | 0.01 | 0.58 | 0.01 |
| WT032 | - | M | 4.8 | 0.52 | 2.09 | 0.25 |
| WT033 | - | M | 7.1 | 0.43 | 1.31 | 0.33 |
| WT034 | - | M | 4.7 | 0 | 3.43 | 0 |
| WT139 | - | M | 5.3 | 0 | 1.59 | 0 |
| WT278 | - | F | 4.8 | 0 | 2.31 | 0 |
| WT330 | - | F | 4.8 | 0 | 1.41 | 0 |
| WT358 | - | F | 5.2 | 0 | 1.20 | 0 |
| WT362 | - | F | 4.3 | 0 | 2.07 | 0 |
| WT403 | - | M | 4.7 | 0 | 2.60 | 0 |
| WT463 | - | M | 4.2 | 0 | 0.85 | 0 |
| WT490 | - | F | 5.5 | 0 | 1.58 | 0 |
| WT_TT | - | F | 5.3 | 0 | 1.57 | 0 |
| *P*-value | | 0.5254 [b] | 0.0127 [c] | 0.0272 [c] | 0.0053 [d] | 0.0009 [d] |

a. ratio of time spent in repetitive circular locomotion to time spent in general locomotion;

b. $\chi^2$ test;

c. independent two sample t-test assuming unequal variance with two tails;

d. independent two sample t-test assuming equal variance with two tails.

### Table S2. Locomotion of 5 TG and 7 WT monkeys in peer separation test with familiar peers

| | ID | Sex | Age (years) | General locomotion (s) | | | Repetitive index in locomotion [a] | | |
|---|---|---|---|---|---|---|---|---|---|
| | | | | Adaptation | Separation | Separation - Adaptation | Adaptation | Separation | Separation - Adaptation |
| TG | TG04 | M | 6.5 | 7 | 340 | 333 | 0 | 0.55 | 0.55 |
| | TG06 | F | 6.5 | 43 | 524 | 481 | 0 | 0.98 | 0.98 |
| | TG08 | M | 6.7 | 196 | 554 | 358 | 0.40 | 0.91 | 0.51 |
| | TG10 | F | 6.5 | 0 | 75 | 75 | 0 | 0 | 0 |
| | TG11 | F | 6.3 | 67 | 265 | 198 | 0 | 0.19 | 0.19 |
| WT | WT002 | M | 7.1 | 125 | 73 | -52 | 0.13 | 0 | -0.13 |
| | WT030 | F | 6.7 | 11 | 221 | 210 | 0 | 0 | 0 |
| | WT032 | M | 6.8 | 69 | 338 | 270 | 0.28 | 0.72 | 0.44 |
| | WT034 | M | 6.8 | 208 | 543 | 335 | 0 | 0 | 0 |
| | WT_12 | F | 14.0 | 47 | 7 | -40 | 0 | 0 | 0 |
| | WT_39 | F | 16.0 | 215 | 327 | 112 | 0 | 0 | 0 |
| | WT_92 | F | 13.0 | 35 | 225 | 190 | 0 | 0 | 0 |
| *P*-value [b] | | 0.9212 [c] | 0.0865 | 0.4375 | 0.3621 | 0.1387 | 0.8048 | 0.0627 | 0.0323 |

a. ratio of time spent in repetitive circular locomotion to time spent in general locomotion;

b. independent two sample *t*-test with two tails;

c. $\chi^2$ test.

### Table S3. Locomotion of 5 TG and 7 WT monkeys in peer separation test with unfamiliar peers

| | ID | Sex | Age (years) | General locomotion (s) | | | Repetitive index in locomotion [a] | | |
|---|---|---|---|---|---|---|---|---|---|
| | | | | Adaptation | Separation | Separation - Adaptation | Adaptation | Separation | Separation - Adaptation |
| TG | TG04 | M | 6.5 | 157 | 42 | -115 | 0.13 | 0.74 | 0.61 |
| | TG06 | F | 6.5 | 152 | 116 | -37 | 0.19 | 0.35 | 0.16 |
| | TG08 | M | 6.7 | 40 | 423 | 383 | 0.73 | 0.89 | 0.15 |
| | TG10 | F | 6.5 | 81 | 309 | 229 | 0 | 0.08 | 0.08 |
| | TG11 | F | 6.3 | 44 | 0 | -44 | 0 | 0 | 0 |
| WT | WT002 | M | 7.1 | 23 | 274 | 252 | 0 | 0.48 | 0.48 |
| | WT030 | F | 6.7 | 0 | 55 | 55 | 0 | 0 | 0 |
| | WT032 | M | 6.8 | 152 | 93 | -59 | 0 | 0.68 | 0.68 |
| | WT034 | M | 6.8 | 133 | 400 | 267 | 0 | 0.16 | 0.16 |
| | WT_12 | F | 14.0 | 47 | 169 | 123 | 0 | 0 | 0 |
| | WT_39 | F | 16.0 | 175 | 338 | 163 | 0 | 0 | 0 |
| | WT_92 | F | 13.0 | 81 | 176 | 96 | 0 | 0 | 0 |
| *P*-value [b] | | 0.9212 [c] | 0.0865 | 0.8389 | 0.6831 | 0.6430 | 0.0905 | 0.2825 | 0.9533 |

a. ratio of time spent in repetitive circular locomotion to time spent in general locomotion;

b. independent two sample *t*-test with two tails;

c. $\chi^2$ test.

### Table S4. EEG data collection from 5 TG and 16 WT monkeys

| ID | Sex | S | Age (years) | Weight (kg) | Heart rate (bpm) | $EtCO_2$ (mmHg) | Temp. (°C) | Isoflurane (%) | N |
|---|---|---|---|---|---|---|---|---|---|
| TG04 | M | 1 | 5.6 | 6.0 | 136 | 27 | 37.6 | 1.40-1.70 | 7 |
| | | 2 | 5.7 | 5.8 | 119 | 27 | 36.2 | 1.30 | 10 |
| | | | **5.6** | **5.9** | | | | | |
| TG06 | F | 1 | 5.5 | 2.8 | 125 | 35 | 36.3 | 1.10-1.20 | 5 |
| | | 2 | 5.7 | 3.1 | 114 | 24 | 35.7 | 1.10-1.20 | 9 |
| | | | **5.6** | **3.0** | | | | | |
| TG08 | M | 1 | 5.6 | 4.9 | 148 | 29 | 37.5 | 1.40-1.50 | 2 |
| | | 2 | 5.7 | 5.1 | 128 | 28 | 35.2 | 1.10-1.50 | 8 |
| | | 3 | 5.8 | 5.2 | 143 | 25 | 36.6 | 1.10-1.60 | 6 |
| | | | **5.7** | **5.1** | | | | | |
| TG10 | F | 1 | 5.6 | 2.8 | 133 | 23 | 35.9 | 1.30 | 6 |
| | | 2 | 5.7 | 2.9 | 117 | 26 | 36.1 | 1.25 | 10 |
| | | | **5.6** | **2.9** | | | | | |
| TG11 | F | 1 | 5.4 | 2.8 | 132 | 24 | 36.9 | 1.20-1.30 | 3 |
| | | 2 | 5.5 | 2.7 | 123 | 29 | 35.5 | 1.10-1.30 | 10 |
| | | | **5.5** | **2.8** | | | | | |
| TG | 2M3F | | 5.60±0.08 [a] | 3.90±1.47 [a] | | | | | 76 |
| WT019 | M | - | 6.5 | 6.8 | 99 | 24 | 35.0 | 0.80-1.00 | 8 |
| WT026 | M | - | 6.3 | 9.6 | 96 | 27 | 36.1 | 1.25 | 6 |
| WT027 | M | - | 6.3 | 9.4 | 102 | 22 | 35.6 | 1.50 | 6 |
| WT030 | F | - | 5.6 | 3.2 | 158 | 26 | 37.2 | 0.70-1.00 | 6 |
| WT032 | M | - | 5.6 | 4.6 | 115 | 25 | 36.1 | 1.30-1.70 | 3 |
| WT033 | M | - | 6.3 | 8.0 | 85 | 28 | 35.8 | 0.80-1.20 | 8 |
| WT034 | M | - | 5.8 | 4.6 | 117 | 29 | 36.9 | 1.20-1.45 | 9 |
| WT139 | M | - | 5.9 | 7.1 | 140 | 27 | 37.1 | 1.25-1.40 | 5 |
| WT278 | F | - | 5.0 | 2.9 | 142 | 24 | 36.7 | 1.30 | 6 |
| WT330 | F | - | 5.1 | 5.2 | 140 | 25 | 37.4 | 1.50 | 6 |
| WT358 | F | - | 4.9 | 4.2 | 126 | 31 | 36.6 | 1.20-1.50 | 8 |
| WT362 | F | - | 5.1 | 2.9 | 150 | 27 | 37.2 | 1.30-1.70 | 6 |
| WT403 | M | - | 5.9 | 6.8 | 125 | 28 | 36.6 | 0.90-1.10 | 8 |
| WT463 | M | - | 5.6 | 6.7 | 124 | 28 | 38.2 | 1.00-1.25 | 3 |
| WT490 | F | - | 4.5 | 3.1 | 130 | 26 | 35.6 | 1.25-1.70 | 7 |
| WT-TT | F | - | 6.2 | 3.1 | 139 | 27 | 37.2 | 1.25-1.50 | 7 |
| WT | 9M7F | | 5.66±0.60 [a] | 5.51±2.30 [a] | | | | | 102 |
| *P*-value | 0.5254 [b] | | 0.8221 [c] | 0.1609 [c] | | | | | |

a. mean ± standard deviation; b. $\chi^2$ test; c. independent two sample *t*-test with two tails.

Bold indicates average values across different sessions from each TG subject.

S, Session; $EtCO_2$, end tidal of $CO_2$; Temp., rectal temperature; M, male; F, female.

**Table S5. Spatial location and anatomical label of all recording electrodes**

| Electrode ID | Electrode label | Full anatomical label |
|---|---|---|
| 2 | OF1 | orbital-frontal, left |
| 6 | OF2 | orbital-frontal, right |
| 3 | PF1 | prefrontal, left |
| 5 | PF2 | prefrontal, right |
| 4 | Fz | frontal, central |
| 9 | FT1 | fronto-temporal, left |
| 16 | FT2 | fronto-temporal, right |
| 19 | T1 | temporal, left |
| 23 | T2 | temporal, right |
| 10 | FP1 | fronto-parietal, left |
| 15 | FP2 | fronto-parietal, right |
| 12 | P1 | medial parietal, left |
| 13 | P2 | medial parietal, right |
| 11 | P3 | lateral parietal, left |
| 14 | P4 | lateral parietal, right |
| 21 | POz | parieto-occipital, central |
| 26 | Oz | occipital, central |
| 25 | O1 | posterior occipital, left |
| 27 | O2 | posterior occipital, right |
| 20 | O3 | anterior occipital, left |
| 22 | O4 | anterior occipital, right |
| | | Electrode location |
| 1 | | top of forehead |
| 7 | | top of nose |
| 8 | | top of left eye |
| 17 | | top of right eye |
| 18 | | horizontally aligned with bottom of left ear |
| 24 | | horizontally aligned with bottom of right ear |

### Table S6. Connections with decreased beta synchronization in TG monkeys

| Number | Electrode1 | Electrode2 | Effect size [a] | *P*-value [b] |
|---|---|---|---|---|
| 1 | Fz | FP1 | -1.0951 | 0.0037 |
| 2 | Fz | PF1 | -1.2140 | 0.0008 |
| 3 | PF1 | OF1 | -1.1328 | 0.0043 |
| 4 | P1 | OF1 | -1.3838 | 0.0015 |
| 5 | P1 | FT1 | -1.3995 | 0.0015 |
| 6 | P1 | FT2 | -1.2004 | 0.0008 |
| 7 | P1 | O2 | -1.4753 | 0.0032 |
| 8 | P1 | O4 | -1.1211 | 0.0018 |
| 9 | P2 | O2 | -1.4020 | 0.0019 |
| 10 | P2 | O4 | -1.0823 | 0.0018 |
| 11 | P4 | O4 | -1.2550 | 0.0006 |

a. *Hedges' g*, TG-WT.

b. independent two sample *t*-test with two tails.

The full anatomical label of electrodes are listed in Table S6.

**Table S7. Cortical and subcortical parcellation and abbreviations**

| Lobes | Number | Hemisphere | Number | Hemisphere | Abbreviation | Full name |
|---|---|---|---|---|---|---|
| Occipital | 1 | L | 2 | R | V1 | Visual area 1 (primary visual cortex) |
| | 3 | L | 4 | R | V2 | Visual area 2 (secondary visual cortex) |
| | 5 | L | 6 | R | VACv | Anterior visual area, ventral part |
| | 7 | L | 8 | R | VACd | Anterior visual area, dorsal part |
| Parietal | 9 | L | 10 | R | S1 | Primary somatosensory cortex |
| | 11 | L | 12 | R | S2 | Secondary somatosensory cortex |
| | 13 | L | 14 | R | PCm | Medial parietal cortex |
| | 15 | L | 16 | R | PCip | Intraparietal cortex |
| | 17 | L | 18 | R | PCi | Inferior parietal cortex |
| | 19 | L | 20 | R | PCs | Superior parietal cortex |
| Temporal | 21 | L | 22 | R | A1 | Primary auditory cortex |
| | 23 | L | 24 | R | A2 | Secondary auditory cortex |
| | 25 | L | 26 | R | TCpol | Temporal polar cortex |
| | 27 | L | 28 | R | TCi | Inferior temporal cortex |
| | 29 | L | 30 | R | TCv | Ventral temporal cortex |
| | 31 | L | 32 | R | TCc | Central temporal cortex |
| | 33 | L | 34 | R | TCs | Superior temporal cortex |
| | 35 | L | 36 | R | HC | Hippocampus |
| | 37 | L | 38 | R | PHC | Parahippocampal cortex |
| PFC | 39 | L | 40 | R | M1 | Primary motor cortex |
| | 41 | L | 42 | R | PMCvl | Ventrolateral premotor cortex |
| | 43 | L | 44 | R | PMCdl | Dorsolateral premotor cortex |
| | 45 | L | 46 | R | PMCm | Medial premotor cortex |
| | 47 | L | 48 | R | FEF | Frontal eye field |
| | 49 | L | 50 | R | PFCvl | Ventrolateral prefrontal cortex |
| | 51 | L | 52 | R | PFCcl | Centrolateral prefrontal cortex |
| | 53 | L | 54 | R | PFCdl | Dorsolateral prefrontal cortex |
| | 55 | L | 56 | R | PFCdm | Dorsomedial prefrontal cortex |
| | 57 | L | 58 | R | PFCm | Medial prefrontal cortex |
| | 59 | L | 60 | R | PFCpol | Prefrontal polar cortex |
| OFC | 61 | L | 62 | R | PFCoi | Orbitoinferior prefrontal cortex |
| | 63 | L | 64 | R | PFCom | Orbitomedial prefrontal cortex |
| | 65 | L | 66 | R | PFCol | Orbitolateral prefrontal cortex |
| Cingulate | 67 | L | 68 | R | CCs | Subgenual cingulate cortex |
| | 69 | L | 70 | R | CCp | Posterior cingulate cortex |
| | 71 | L | 72 | R | CCr | Retrosplenial cingulate cortex |
| | 73 | L | 74 | R | CCa | Anterior cingulate cortex |

| | | | | | | |
|---|---|---|---|---|---|---|
| Insula | 75 | L | 76 | R | G | Gustatory cortex |
| | 77 | L | 78 | R | Ia | Anterior insula |
| | 79 | L | 80 | R | Ip | Posterior insula |
| Subcortical | 81 | L | 82 | R | Amyg | Amygdala |
| | 83 | L | 84 | R | Cau | Caudate |
| | 85 | L | 86 | R | Put | Putamen |
| | 87 | L | 88 | R | Tha | Thalamus |
| | 89 | L | 90 | R | HT | Hypothalamus |
| | 91 | L | 92 | R | NAcc | Nucleus accumbens |
| | 93 | L | 94 | R | GP | Globus pallidus |

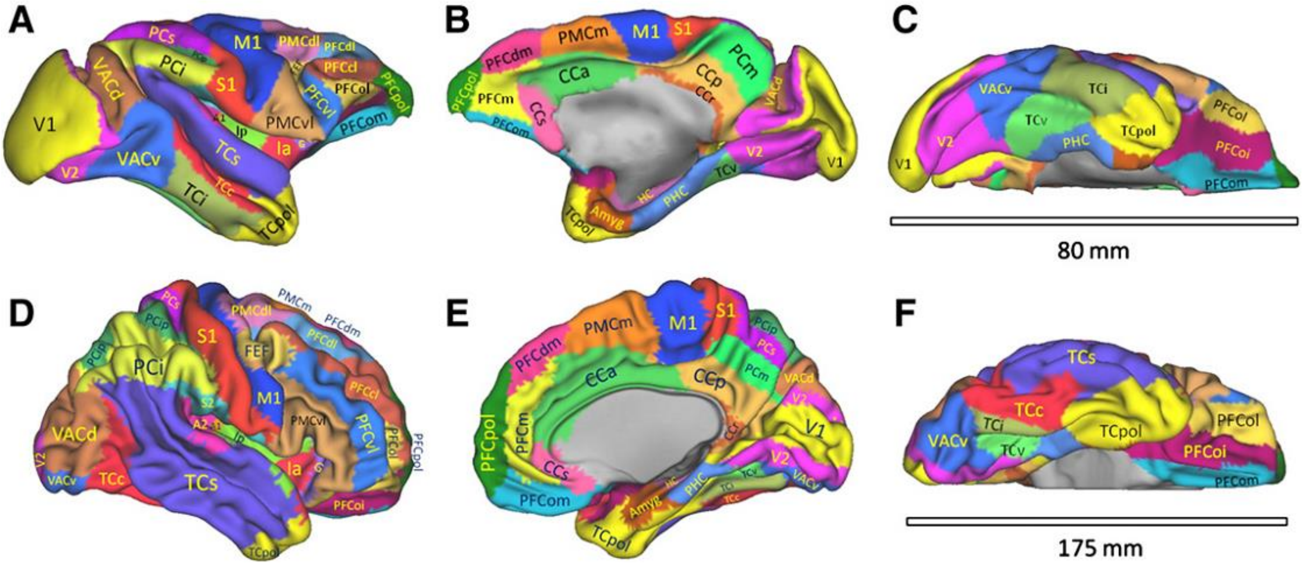


Regional Map representations on macaque F99 (A, B, C) and human MNI (D, E, F) templates. The figure is adopted, with permission, from REF. 36.

**Table S8. MRI data collection from 5 TG and 11 WT monkeys**

| ID | Gender | Copy number | Weight (kg) | Age (year) | Heart rate (beat/min) | EtCO$_2$ (mmHg) | Temp. (°C) | Isoflurane (%) | Run |
|---|---|---|---|---|---|---|---|---|---|
| TG04 | M | 1.0 | 4.3 | 4.9 | ~115 | ~29 | ~37.0 | 1.0 | 9 |
| TG06 | F | 7.3 | 2.7 | 4.3 | ~103 | ~24 | ~35.6 | 1.0 | 8 |
| TG08 | M | 2.9 | 3.8 | 4.2 | ~125 | ~29 | ~36.1 | 1.0 | 7 |
| TG10 | F | 1.1 | 2.6 | 4.2 | ~94 | ~26 | ~35.8 | 1.0 | 8 |
| TG11 | F | 1.9 | 2.9 | 4.4 | ~130 | ~27 | ~36.8 | 1.2 | 13 |
| TG | 2M3F | | 3.26±0.75[a] | 4.40±0.29 [a] | | | | | 45 |
| WT030 | F | - | 3.5 | 4.8 | ~148 | ~27 | ~37.7 | 0.8 | 8 |
| WT032 | M | - | 3.9 | 4.4 | ~120 | ~30 | ~37.7 | 1.0 | 10 |
| WT034 | M | - | 3.5 | 4.4 | ~120 | ~30 | ~37.0 | 1.0 | 7 |
| WT139 | M | - | 7 | 5.5 | ~148 | ~27 | ~37.2 | 1.2 | 10 |
| WT278 | F | - | 3.4 | 4.5 | ~125 | ~28 | ~35.6 | 1.2-1.25 | 10 |
| WT330 | F | - | 4.3 | 4.6 | ~125 | ~27 | ~35.9 | 1.2-1.3 | 10 |
| WT358 | F | - | 3.2 | 4.2 | ~130 | ~28 | ~37.8 | 1.25-1.3 | 10 |
| WT362 | F | - | 3 | 4.5 | ~130 | ~27 | ~36.1 | 1.2 | 10 |
| WT463 | M | - | 6.1 | 5.1 | ~110 | ~27 | ~37.2 | 1.3-1.2 | 5 |
| WT490 | F | - | 2.9 | 4.1 | ~136 | ~30 | ~37.1 | 1.1 | 9 |
| WT_TT | F | - | 2.9 | 5.4 | ~159 | ~27 | ~37.0 | 1.2 | 10 |
| WT | 4M7F | | 3.97±1.36[a] | 4.68±0.46 [a] | | | | | 99 |
| *P*-value | 0.89 [b] | | | 0.233 [c] | | | | | |

Abbreviations. EtCO$_2$, end tidal of CO$_2$. Temp., rectal temperature.

a. mean ± SD; b. $\chi^2$ test; c. two sample Student's *t*-test with two tails

**Table S9. Abnormal functional connections in transgenic monkeys**

| Number | Region1 | Region2 | Effect size | -log(p) | Number | Region1 | Region2 | Effect size | -log(p) |
|---|---|---|---|---|---|---|---|---|---|
| 1 | PFCol.R | TCpol.L | 3.7934 | 3.0883 | 26 | CCa.L | CCp.R | 1.9535 | 3.9254 |
| 2 | S1.R | TCpol.L | -2.1053 | 3.8017 | 27 | PFCpol.L | CCr.L | 2.2171 | 3.1671 |
| 3 | PFCoi.L | Amyg.L | 2.1135 | 3.3893 | 28 | CCa.L | CCr.L | 1.5189 | 3.0292 |
| 4 | PFCol.R | Amyg.L | 1.7148 | 3.1742 | 29 | PFCcl.R | CCr.L | 2.5315 | 3.4020 |
| 5 | PFCom.L | PHC.L | 1.9072 | 3.0448 | 30 | PFCpol.L | CCr.R | 2.5455 | 3.4546 |
| 6 | FEF.R | PHC.L | 2.5356 | 3.7297 | 31 | CCa.L | CCr.R | 1.8818 | 3.7185 |
| 7 | PFCcl.R | PHC.L | 3.3543 | 4.5150 | 32 | PFCcl.R | CCr.R | 3.1228 | 3.2941 |
| 8 | PMCdl.R | PHC.L | 2.2256 | 3.2446 | 33 | PCi.L | PFCol.R | 1.6494 | 3.3683 |
| 9 | S1.L | TCi.L | -2.0907 | 3.3084 | 34 | PCi.R | PFCol.R | 2.0144 | 3.7815 |
| 10 | S1.R | TCi.L | -1.7644 | 3.4017 | 35 | PCm.R | PFCvl.R | 2.0484 | 3.7557 |
| 11 | CCa.L | TCv.L | 1.9641 | 3.0130 | 36 | PCm.R | PFCpol.L | 2.3013 | 3.0973 |
| 12 | FEF.R | HC.L | 2.0275 | 3.6536 | 37 | FEF.L | PFCpol.L | 1.5896 | 3.1402 |
| 13 | S1.R | HC.R | -3.8493 | 4.6105 | 38 | FEF.R | PFCpol.L | 1.8318 | 3.7211 |
| 14 | CCa.L | HC.R | 2.0348 | 3.4741 | 39 | PFCcl.R | PFCpol.L | 2.0013 | 3.0732 |
| 15 | PFCcl.L | HC.R | 1.9149 | 3.5901 | 40 | PCi.L | PFCpol.L | 1.6887 | 3.1420 |
| 16 | PFCol.R | TCc.L | 2.3282 | 4.5556 | 41 | PCs.R | PFCpol.L | 2.0421 | 3.8437 |
| 17 | S1.R | TCc.L | -3.8307 | 5.8539 | 42 | PFCdm.R | PFCpol.L | 2.2320 | 3.0922 |
| 18 | PFCol.R | TCc.R | 2.1251 | 3.5383 | 43 | PFCdl.L | PFCpol.L | 2.1559 | 3.5973 |
| 19 | S1.R | V1.R | -2.2162 | 4.4059 | 44 | PFCdl.R | PFCpol.L | 2.9842 | 3.8474 |
| 20 | Ip.R | PFCoi.R | 1.8808 | 3.7257 | 45 | PFCdl.R | S1.R | -1.4825 | 3.0640 |
| 21 | PCm.R | PFCoi.R | 1.7902 | 3.5763 | 46 | PFCcl.R | PFCm.R | 1.8257 | 3.0204 |
| 22 | CCr.L | PFCom.R | 1.7261 | 3.1949 | 47 | PFCdl.R | PFCm.R | 1.5808 | 3.2776 |
| 23 | PFCdl.L | PFCom.R | 1.591 | 3.0762 | 48 | CCa.L | PCm.L | 1.8289 | 3.2516 |
| 24 | PFCol.L | Ip.R | 2.4204 | 3.2313 | 49 | CCa.L | PCm.R | 2.4953 | 4.9584 |
| 25 | CCa.L | CCp.L | 1.8406 | 3.5798 | 50 | PFCdm.R | PCs.R | 2.0269 | 3.4534 |

### Table S10. Group different edges correlating with behavioral measures

| Region1 | Region2 | Spearman's *r* | edge-wise *P* |
|---|---|---|---|
| Egdes correlated with general locomotion | | | |
| S1.R | TCi.L | -0.79 | 0.0005 |
| S1.R | HC.R | -0.79 | 0.0005 |
| S1.R | TCc.L | -0.74 | 0.0014 |
| PFCdl.R | S1.R | -0.70 | 0.0032 |
| S1.L | TCi.L | -0.70 | 0.0034 |
| PFCol.L | Ip.R | 0.66 | 0.0065 |
| S1.R | V1.R | -0.66 | 0.0072 |
| Egdes correlated with regressive error rate | | | |
| PFCpol.L | CCr.L | 0.95 | 0.0004 |
| PMCdl.R | PHC.L | 0.90 | 0.0020 |
| PFCom.L | PHC.L | 0.88 | 0.0031 |
| PFCcl.R | PHC.L | 0.83 | 0.0083 |
| PFCpol.L | CCr.R | 0.82 | 0.0108 |
| PFCdm.R | PFCpol.L | 0.82 | 0.0108 |
| PFCdl.L | PFCpol.L | 0.82 | 0.0108 |
| PFCdl.R | PFCpol.L | 0.78 | 0.0172 |
| PFCdl.L | PFCom.R | 0.75 | 0.0255 |
| PFCdl.R | PFCm.R | 0.75 | 0.0255 |
| PFCcl.R | PFCpol.L | 0.73 | 0.0311 |
| CCa.L | HC.R | 0.72 | 0.0369 |
| PCi.L | PFCpol.L | 0.72 | 0.0369 |
| CCa.L | CCr.L | 0.70 | 0.0433 |

**Table S11. Characteristics of all human participants**

| | Autism | | | Autism with ADOS | | | | | TDC | | |
|---|---|---|---|---|---|---|---|---|---|---|---|
| Site | Number (M/F) | Age,mean(SD) [range] | FIQ,mean(SD) [range] | Number (M/F) | ADOS_TOTAL mean(SD) [range] | ADOS_COMM mean(SD) [range] | ADOS_SOCIAL mean(SD) [range] | ADOS_STEREO mean(SD) [range] | Number (M/F) | Age,mean (SD) [range] | FIQ,mean (SD) [range] |
| NYU | 9(8/1) | 13.78(1.52) | 101.33(11.98) | 9(8/1) | 12.67(4.77) | 4.00(1.50) | 8.67(3.54) | 3.33(1.66) | 35(26/9) | 14.65(1.72) | 109.74(13.93) |
| | | [12.37-16.74] | [90-128] | | [5-19] | [2-6] | [2-13] | [2-7] | | [12.10-17.70] | [81-126] |
| PITT | 12(9/3) | 14.09(1.93) | 109.17(13.20) | 9(7/2) | 11.67(3.39) | 3.67(1.22) | 8.00(2.45) | 1.78(1.20) | 12(9/3) | 14.59(1.71) | 107.33(8.97) |
| | | [12.03-17.78] | [87-126] | | [7-17] | [2-5] | [5-12] | [0-3] | | [12.15-17.36] | [97-130] |
| TRINITY | 8(8/0) | 14.91(1.89) | 103.62(13.05) | -- | -- | -- | -- | -- | 14(14/0) | 14.58(1.73) | 106.43(12.33) |
| | | [12.75-17.25] | [89-122] | | -- | -- | -- | -- | | [12.04-17.50] | [89-128] |
| UCLA_1 | 20(18/2) | 14.77(1.73) | 104.85(10.64) | 20(18/2) | 9.00(3.91) | 2.90(1.52) | 6.10(2.77) | 0.85(1.27) | 20(17/3) | 14.19(1.66) | 102.75(9.71) |
| | | [12.33-17.94] | [86-127] | | [2-16] | [0-5] | [2-11] | [0-5] | | [12.01-17.79] | [84-126] |
| UCLA_2 | 5(5/0) | 14.08(1.53) | 95.00(7.68) | 4(4/0) | 12.75(3.30) | 3.25(0.96) | 9.50(2.38) | 3.00(2.45) | 5(3/2) | 12.96(0.59) | 111.80(11.84) |
| | | [12.82-16.47] | [86-104] | | [9-16] | [2-4] | [7-12] | [1-6] | | [12.15-13.63] | [99-128] |

**Table S11. Continued**

| | Autism | | | Autism with ADOS | | | | | TDC | | |
|---|---|---|---|---|---|---|---|---|---|---|---|
| Site | Number (M/F) | Age,mean(SD) [range] | FIQ,mean(SD) [range] | Number (M/F) | ADOS_TOTAL mean(SD) [range] | ADOS_COMM mean(SD) [range] | ADOS_SOCIAL mean(SD) [range] | ADOS_STEREO mean(SD) [range] | Number (M/F) | Age,mean (SD) [range] | FIQ,mean (SD) [range] |
| UM_1 | 14(11/3) | 14.09(1.59) | 105.57(11.55) | -- | -- | -- | -- | -- | 22(13/9) | 15.00(1.89) | 107.75(10.03) |
| | | [12.40-18.00] | [89.50-126] | | -- | -- | -- | -- | | [12.20-17.90] | [89-124.50] |
| UM_2 | 7(6/1) | 15.46(1.24) | 109.86(14.35) | -- | -- | -- | -- | -- | 17(16/1) | 15.59(1.46) | 109.97(10.15) |
| | | [13.10-16.60] | [90.50-129.50] | | -- | -- | -- | -- | | [13.60-17.90] | [89.50-129] |
| USM | 13(13/0) | 16.39(1.54) | 97.46(12.69) | 12(12/0) | 13.42(3.00) | 4.33(1.67) | 9.08(1.83) | 2.17(1.03) | 5(5/0) | 14.99(1.67) | 114.60(15.16) |
| | | [12.33-17.79] | [83-118] | | [7-18] | [1-7] | [6-12] | [0-4] | | [12.94-17.07] | [97-130] |
| YALE | 2(0/2) | 15.46(1.47) | 86.50(2.12) | -- | -- | -- | -- | -- | 10(6/4) | 14.45(1.25) | 102.40(10.36) |
| | | [14.42-16.50] | [85-88] | | -- | -- | -- | -- | | [12.75-16.66] | [89-120] |
| SUM | 90(78/12) | 14.75(1.79) | 103.44(12.44) | 54(49/5) | 11.31(4.08) | 3.56(1.54) | 7.76(2.89) | 1.87(1.63) | 140(109/31) | 14.68(1.69) | 107.64(11.63) |
| | | [12.03-18] | [83-129.50] | | [2-19] | [0-7] | [2-13] | [0-7] | | [12.01-17.90] | [81-130] |
| *P* value | 0.0944 [a] | 0.7564 [b] | 0.0099 [b] | | | | | | | | |

Abbreviations. TDC, typically developing control; ADOS, Autism Diagnostic Observation Schedule; ADOS_TOTAL, classic total ADOS score; ADOS_COMM, communication total sub-score of the classic ADOS; ADOS_SOCIAL, social total score of the classic ADOS; ADOS_STEREO, stereotyped behaviors and restricted interests total sub-score of the classic ADOS.

a. $\chi^2$ test; b. two sample Student's *t*-test with two tails. "--" indicates there are no participants with clinical scores of ADOS.

### Table S12. Characteristics of human participants in subgroup 1

| | Autism | | | Autism with ADOS | | | | | TDC | | |
|---|---|---|---|---|---|---|---|---|---|---|---|
| Site | Number (M/F) | Age,mean(SD) [range] | FIQ,mean(SD) [range] | Number (M/F) | ADOS_TOTAL mean(SD) [range] | ADOS_COMM mean(SD) [range] | ADOS_SOCIAL mean(SD) [range] | ADOS_STEREO mean(SD) [range] | Number (M/F) | Age,mean (SD) [range] | FIQ,mean (SD) [range] |
| NYU | 5(4/1) | 14.23(1.84) [12.64-16.74] | 97.00(8.22) [90-109] | 5(4/1) | 13.60(5.18) [5-19] | 4.40(1.34) [3-6] | 9.20(4.21) [2-13] | 4.00(1.87) [2-7] | 15(11/4) | 14.65(1.76) [12.10-17.31] | 110.00(14.36) [81-126] |
| PITT | 7(5/2) | 13.33(0.79) [12.24-14.20] | 106.86(12.29) [96-126] | 4(3/1) | 11.75(3.30) [8-16] | 3.50(1.29) [2-5] | 8.25(2.50) [5-11] | 1.50(1.29) [0-3] | 7(5/2) | 14.16(1.27) [12.83-15.82] | 105.71(6.32) [97-114] |
| TRINITY | 2(2/0) | 15.43(2.26) [13.83-17.03] | 97.50(0.71) [97-98] | -- | -- -- | -- -- | -- -- | -- -- | 9(9/0) | 14.85(1.76) [12.04-17.50] | 105.44(12.97) [89-128] |
| UCLA_1 | 12(11/1) | 14.82(1.92) [12.33-17.53] | 105.58(13.37) [86-127] | 12(11/1) | 9.58(3.82) [3-16] | 3.17(1.59) [1-5] | 6.42(2.50) [2-11] | 1.17(1.47) [0-5] | 12(10/2) | 14.09(1.59) [12.01-17.79] | 105.00(10.44) [84-126] |
| UCLA_2 | 2(2/0) | 14.77(2.40) [13.08-16.47] | 89.00(4.24) [86-92] | 2(2/0) | 13.50(3.54) [11-16] | 3.50(0.71) [3-4] | 10.00(2.83) [8-12] | 3.50(3.54) [1-6] | 3(1/2) | 13.00(0.36) [12.64-13.36] | 104.33(6.81) [99-112] |

### Table S12. Continued

| | Autism | | | Autism with ADOS | | | | | TDC | | |
|---|---|---|---|---|---|---|---|---|---|---|---|
| Site | Number (M/F) | Age,mean(SD) [range] | FIQ,mean(SD) [range] | Number (M/F) | ADOS_TOTAL mean(SD) [range] | ADOS_COMM mean(SD) [range] | ADOS_SOCIAL mean(SD) [range] | ADOS_STEREO mean(SD) [range] | Number (M/F) | Age,mean (SD) [range] | FIQ,mean (SD) [range] |
| UM_1 | 8(7/1) | 14.64(1.79) | 107.62(10.61) | -- | -- | -- | -- | -- | 10(6/4) | 15.17(1.76) | 109.45(8.18) |
| | | [12.80-18] | [96-124.50] | | -- | -- | -- | -- | | [12.40-17.80] | [96.50-124.50] |
| UM_2 | 5(4/1) | 15.38(1.47) | 113.60(14.05) | -- | -- | -- | -- | -- | 9(8/1) | 15.83(1.67) | 111.00(12.55) |
| | | [13.10-16.60] | [94-129.50] | | -- | -- | -- | -- | | [13.60-17.90] | [89.50-129] |
| USM | 7(7/0) | 16.20(1.80) | 89.43(8.66) | 6(6/0) | 13.83(2.23) | 4.33(1.21) | 9.50(1.87) | 2.00(0.63) | 2(2/0) | 13.37(0.62) | 128.00(2.83) |
| | | [12.33-17.65] | [83-107] | | [10-16] | [3-6] | [7-12] | [1-3] | | [12.94-13.81] | [126-130] |
| YALE | 1(0/1) | 14.42(--) | 85.00(--) | -- | -- | -- | -- | -- | 5(4/1) | 14.62(1.15) | 100.80(7.60) |
| | | [14.42-14.42] | [85-85] | | -- | -- | -- | -- | | [13.33-15.92] | [92-110] |
| SUM | 49(42/7) | 14.79(1.78) | 102.31(13.17) | 29(26/3) | 11.72(3.98) | 3.69(1.42) | 8.03(2.95) | 2.03(1.80) | 72(56/16) | 14.65(1.65) | 107.85(11.46) |
| | | [12.24-18] | [83-129.50] | | [3-19] | [1-6] | [2-13] | [0-7] | | [12.01-17.90] | [81-130] |
| *P* | 0.2747 [a] | 0.6612 [b] | 0.0154 [b] | | | | | | | | |

Abbreviations. TDC, typically developing control; ADOS, Autism Diagnostic Observation Schedule; ADOS_TOTAL, classic total ADOS score; ADOS_COMM, communication total sub-score of the classic ADOS; ADOS_SOCIAL, social total score of the classic ADOS; ADOS_STEREO, stereotyped behaviors and restricted interests total sub-score of the classic ADOS.

a. $\chi^2$ test; b. two sample Student's *t*-test with two tails; "--" indicates there are no participants with clinical scores of ADOS.

### Table S13. Characteristics of human participants in subgroup 2

| | Autism | | | Autism with ADOS | | | | | TDC | | |
|---|---|---|---|---|---|---|---|---|---|---|---|
| Site | Number (M/F) | Age,mean(SD) [range] | FIQ,mean(SD) [range] | Number (M/F) | ADOS_TOTAL mean(SD) [range] | ADOS_COMM mean(SD) [range] | ADOS_SOCIAL mean(SD) [range] | ADOS_STEREO mean(SD) [range] | Number (M/F) | Age,mean (SD) [range] | FIQ,mean (SD) [range] |
| NYU | 4(4/0) | 13.21(0.94) [12.37-14.53] | 106.75(14.91) [95-128] | 4(4/0) | 11.50(4.65) [7-18] | 3.50(1.73) [2-6] | 8.00(2.94) [5-12] | 2.50(1.00) [2-4] | 20(15/5) | 14.64(1.73) [12.10-17.70] | 109.55(13.97) [83-126] |
| PITT | 5(4/1) | 15.14(2.63) [12.03-17.78] | 112.40(15.18) [87-124] | 5(4/1) | 11.60(3.85) [7-17] | 3.80(1.30) [2-5] | 7.80(2.68) [5-12] | 2.00(1.22) [0-3] | 5(4/1) | 15.19(2.20) [12.15-17.36] | 109.60(12.26) [98-130] |
| TRINITY | 6(6/0) | 14.73(1.96) [12.75-17.25] | 105.67(14.77) [89-122] | -- | -- -- | -- -- | -- -- | -- -- | 5(5/0) | 14.10(1.74) [12.25-15.91] | 108.20(12.32) [97-126] |
| UCLA_1 | 8(7/1) | 14.69(1.54) [13.37-17.94] | 103.75(4.92) [98-112] | 8(7/1) | 8.12(4.12) [2-14] | 2.50(1.41) [0-5] | 5.62(3.25) [2-10] | 0.38(0.74) [0-2] | 8(7/1) | 14.33(1.85) [12.36-17.78] | 99.38(7.93) [88-110] |
| UCLA_2 | 3(3/0) | 13.62(1.02) [12.82-14.77] | 99.00(7.00) [91-104] | 2(2/0) | 12.00(4.24) [9-15] | 3.00(1.41) [2-4] | 9.00(2.83) [7-11] | 2.50(2.12) [1-4] | 2(2/0) | 12.89(1.05) [12.15-13.63] | 123.00(7.07) [118-128] |

### Table S13. Continued

| | Autism | | | Autism with ADOS | | | | | TDC | | |
|---|---|---|---|---|---|---|---|---|---|---|---|
| Site | Number (M/F) | Age,mean(SD) [range] | FIQ,mean(SD) [range] | Number (M/F) | ADOS_TOTAL mean(SD) [range] | ADOS_COMM mean(SD) [range] | ADOS_SOCIAL mean(SD) [range] | ADOS_STEREO mean(SD) [range] | Number (M/F) | Age,mean (SD) [range] | FIQ,mean (SD) [range] |
| UM_1 | 6(4/2) | 13.37(0.99) | 102.83(13.17) | -- | -- | -- | -- | -- | 12(7/5) | 14.87(2.05) | 106.33(11.51) |
| | | [12.40-15.00] | [89.50-126] | | -- | -- | -- | -- | | [12.20-17.90] | [89-122] |
| UM_2 | 2(2/0) | 15.65(0.64) | 100.50(14.14) | -- | -- | -- | -- | -- | 8(8/0) | 15.31(1.25) | 108.81(7.26) |
| | | [15.20-16.10] | [90.50-110.50] | | -- | -- | -- | -- | | [14-17.60] | [93.50-116] |
| USM | 6(6/0) | 16.60(1.30) | 106.83(10.03) | 6(6/0) | 13.00(3.79) | 4.33(2.16) | 8.67(1.86) | 2.33(1.37) | 3(3/0) | 16.07(1.02) | 105.67(12.50) |
| | | [14.14-17.79] | [95-118] | | [7-18] | [1-7] | [6-11] | [0-4] | | [15.03-17.07] | [97-120] |
| YALE | 1(0/1) | 16.50(--) | 88.00(--) | -- | -- | -- | -- | -- | 5(2/3) | 14.28(1.45) | 104.00(13.32) |
| | | [16.50-16.50] | [88-88] | | -- | -- | -- | -- | | [12.75-16.66] | [89-120] |
| SUM | 41(36/5) | 14.71(1.82) | 104.80(11.51) | 25(23/2) | 10.84(4.22) | 3.40(1.68) | 7.44(2.86) | 1.68(1.41) | 68(53/15) | 14.71(1.73) | 107.42(11.89) |
| | | [12.03-17.94] | [87-128] | | [2-18] | [0-7] | [2-12] | [0-4] | | [12.10-17.90] | [83-130] |
| *p* | 0.1975 [a] | 0.9937 [b] | 0.2631[b] | | | | | | | | |

Abbreviations. TDC, typically developing control; ADOS, Autism Diagnostic Observation Schedule; ADOS_TOTAL, classic total ADOS score; ADOS_COMM, communication total sub-score of the classic ADOS; ADOS_SOCIAL, social total score of the classic ADOS; ADOS_STEREO, stereotyped behaviors and restricted interests total sub-score of the classic ADOS.

a. $\chi^2$ test; b. two sample Student's *t*-test with two tails; "--" indicates there are no participants with clinical scores of ADOS.

### Table S14. Disrupted functional connections in human autism of subgroup 1

| Number | Region1 | Region2 | Effect size | -log($P$) | Number | Region1 | Region2 | Effect size | -log($P$) |
|---|---|---|---|---|---|---|---|---|---|
| 1 | TCv.R | TCpol.L | -0.7020 | 3.7249 | 17 | PFCpol.R | TCv.R | -0.6270 | 3.2901 |
| 2 | PHC.L | TCpol.R | -0.7269 | 3.7813 | 18 | PFCol.L | TCs.L | -0.7141 | 3.2123 |
| 3 | PHC.R | TCpol.R | -0.6503 | 3.4031 | 19 | V1.L | TCs.R | -0.6740 | 3.3610 |
| 4 | TCi.L | TCpol.R | -0.6777 | 3.5117 | 20 | V1.R | TCs.R | -0.6651 | 3.2372 |
| 5 | TCv.L | TCpol.R | -0.7364 | 3.9847 | 21 | V2.L | TCs.R | -0.7216 | 3.7954 |
| 6 | TCv.R | TCpol.R | -0.8461 | 5.0609 | 22 | A1.R | V1.L | -0.6562 | 3.2660 |
| 7 | V2.L | TCpol.R | -0.6908 | 3.4606 | 23 | A1.R | V1.R | -0.6983 | 3.7038 |
| 8 | PFCol.L | TCpol.R | -0.7289 | 3.4295 | 24 | A1.R | V2.R | -0.6556 | 3.3549 |
| 9 | PFCcl.L | TCpol.R | -0.7048 | 3.1574 | 25 | PFCol.R | Ia.R | -0.6785 | 3.1840 |
| 10 | TCv.L | Amyg.L | -0.6464 | 3.0862 | 26 | PFCol.R | Ip.L | -0.7350 | 3.7958 |
| 11 | TCv.R | Amyg.L | -0.6971 | 3.5103 | 27 | PFCvl.R | Ip.L | -0.7080 | 3.5224 |
| 12 | TCs.R | TCi.L | -0.7266 | 3.7823 | 28 | PFCpol.R | CCp.L | -0.7409 | 3.7102 |
| 13 | TCs.R | TCi.R | -0.6976 | 3.4891 | 29 | PFCpol.R | CCr.L | -0.6533 | 3.1049 |
| 14 | TCs.R | TCv.L | -0.7629 | 4.1828 | 30 | A1.L | PFCol.R | -0.6591 | 3.3045 |
| 15 | PFCpol.R | TCv.L | -0.6236 | 3.1153 | 31 | A1.R | PFCol.R | -0.6620 | 3.3096 |
| 16 | TCs.R | TCv.R | -0.8382 | 4.7878 | 32 | PCi.R | A1.L | -0.6364 | 3.0604 |

**Table S15. Correlations between connectivity fingerprint of human autism in subgroup 1 and symptom severity**

| | TOTAL | COMM | SOCIAL | STEREO |
|---|---|---|---|---|
| Spearman's *r* | 0.2427 | **0.4162** | 0.1796 | 0.0361 |
| *P* | 0.2046 | **0.0247** | 0.3511 | 0.8525 |
| Pearson's *r* | 0.2709 | 0.3507 | 0.1974 | -0.0094 |
| *P* | 0.1552 | 0.0622* | 0.3046 | 0.9613 |

TOTAL, classic total ADOS score; COMM, communication total sub-score of the classic ADOS; SOCIAL, social total score of the classic ADOS; STEREO, stereotypic behaviors and restricted interests total sub-score of the classic ADOS.

*marginally significant.

**Table S16. Correlations of group differences in connectivity networks between monkeys and humans (subgroup 1)**

| | Occipital | Parietal | Temporal | PFC | OFC | Cingulate | Insula | Subcortical |
|---|---|---|---|---|---|---|---|---|
| Occipital | 0.1404 | - | - | - | - | - | - | - |
| *P* value | 0.4759 | - | - | - | - | - | - | - |
| Parietal | 0.0564 | 0.2923 | - | - | - | - | - | - |
| *P* value | 0.5854 | 0.0173 | - | - | - | - | - | - |
| Temporal | **0.5614** | **-0.3127** | -0.0078 | - | - | - | - | - |
| *P* value | **2.49e-13** | **2.76e-6** | 0.924 | - | - | - | - | - |
| PFC | 0.0198 | **-0.2246** | **-0.3158** | 0.0525 | - | - | - | - |
| *P* value | 0.7944 | **0.0002** | **1.27e-10** | 0.4272 | - | - | - | - |
| OFC | 0.1869 | 0.1824 | -0.1258 | 0.0362 | 0.2463 | - | - | - |
| *P* value | 0.2033 | 0.1251 | 0.1947 | 0.6802 | 0.3761 | - | - | - |
| Cingulate | 0.1744 | -0.1059 | **0.3976** | **-0.3749** | -0.0741 | 0.2966 | - | - |
| *P* value | 0.1680 | 0.3044 | **8.02e-7** | **2.94e-7** | 0.6167 | 0.1253 | - | - |
| Insula | **0.5083** | -0.1161 | 0.1948 | **-0.3566** | 0.0025 | -0.2361 | -0.0994 | - |
| *P* value | **0.0002** | 0.3314 | 0.0433 | **2.70e-5** | 0.9883 | 0.1062 | 0.7246 | - |
| Subcortical | 0.1116 | -0.1223 | **-0.4044** | **-0.2547** | 0.189 | -0.2674 | -0.2309 | 0.0033 |
| *P* value | 0.2414 | 0.1143 | **2.46e-11** | **5.98e-6** | 0.0852 | 0.0044 | 0.0346 | 0.9749 |

Bold font, statistically significant correlation after Bonferroni correction for multiple comparisons.
